# Genome Assembly of the Iconic Samba Mahsuri Delineates Locus-specific Population Structure within *Indica* Rice

**DOI:** 10.64898/2026.04.20.719576

**Authors:** Deepti Rao, Shashi Kiran K., Nikhila Tummala, Erum Khan, Ramesh V. Sonti, Shrish Tiwari, Hitendra K. Patel

## Abstract

High-quality reference genomes enable detailed analysis of structural variation and its consequences for genome organization in crops. Here, we present a chromosome-scale genome assembly of *Oryza sativa* cv. Samba Mahsuri (SM), an elite Indian mega rice variety cultivated for its grain and cooking quality. Using PacBio HiFi sequencing in combination with Illumina reads and Bionano optical mapping, we generated a ∼395 Mb assembly (SMv1.0) with 97.7% BUSCO completeness. A robust annotation framework identified 31,138 evidence-guided protein-coding gene models alongside 59,152 ab initio predictions. Comparative genomic analyses revealed extensive macrosynteny with established rice reference genomes, while uncovering pronounced locus-specific sequence and structural polymorphisms. Notably, a complex inversion-match-inversion (IMI) configuration on chromosome 6 differentiates SM from the *japonica* reference Nipponbare, but not from the *indica* reference R498. Population-scale analyses of 533 cultivated and 4 wild rice accessions demonstrate that genetic variation within the IMI region produces a markedly sharper and more coherent population structure than is observed in flanking regions or genome-wide, including tight subpopulation-based clustering and segregation of alternative IMI configurations within *indica* rice. Together, these results establish SMv1.0 as a robust chromosome-scale reference genome sequence for rice and demonstrate how large structural polymorphisms can shape locus-specific patterns of relatedness that diverge from genome-wide ancestry.

**Significance Statement:** We present a high-quality chromosome-scale genome sequence of the elite Indian rice variety Samba Mahsuri (SM), which, to the best of our knowledge, represents the first chromosome-scale reference genome from Indian rice germplasm assembled using a map-based method. Using this reference to analyze population-scale genotyping data from 533 cultivated and four wild rice accessions reveals markedly tighter population clustering within a megabase-scale inversion (IMI) region, than at the whole-genome scale, along with a pronounced split within *indica* rice that is independent of genome-wide ancestry.

## Introduction

Rice (*Oryza sativa* L.) is one of the world’s most important staple crops and a central model for studying genome evolution and genomic variation (Molina et al., 2011; Qin et al., 2021; Xie et al., 2015). Rice harbors extensive genetic diversity preserved across global germplasm collections, yet reference-quality genomes remain biased toward a limited set of cultivars. As one of the world’s largest producers of rice, India maintains over one lakh germplasm accessions through the National Gene Bank and associated repositories, representing a substantial but under-characterized component of this diversity (https://icar-crri.in, https://www.pib.gov.in). Here, we present Samba Mahsuri as a map-based reference-grade genome to bridge this gap.

Samba Mahsuri (BPT 5204; hereafter referred to as SM) is a widely cultivated elite Indian mega rice variety valued for its high yield, semi-dwarf architecture, and superior grain quality. Developed through crosses involving elite *indica* cultivars and an indica-japonica hybrid (GEB-24 × TN(1)/Mahsuri), SM represents an improved indica genetic background with limited japonica introgression (Reddy et al., 1979). Owing to its agronomic performance and grain quality, SM is extensively grown in India and is frequently used in breeding programs (Potupureddi et al., 2021; Subhra Priyadarshini, 2025; Sundaram et al., 2008). A previous genomic analysis of SM relied on short-read resequencing against the Nipponbare reference to identify SNPs and short INDELs associated with grain quality and stress-related traits (Reddy Lachagari et al., 2019). However, the absence of a chromosome-scale reference genome has limited the characterization of larger-scale genomic variation in this cultivar.

Structural variation, including large insertions, deletions, duplications, and inversions, represents a major source of genomic diversity in plant genomes (Jayakodi et al., 2020; W. Wang et al., 2018). In particular, chromosomal inversions can suppress recombination across extended genomic intervals, facilitating the persistence of divergent haplotypes and influencing patterns of genetic diversity, linkage disequilibrium, and population structure, infrequently giving rise to more drastic effects, like speciation (Crow et al., 2020; Lundberg et al., 2023; Mathiopoulos & Lanzaro, 1995; Szamalek et al., 2006). A recent rice pan-genome study has revealed that inversions are widespread across Asian rice, often segregating within subpopulations rather than being fixed within subspecies, and in some cases predating domestication (Zhou et al., 2023). These findings underscore the importance of structural variation in shaping genome evolution. Because population-scale analyses are inherently reference-dependent, high-quality genome assemblies from diverse genetic backgrounds are essential for resolving locus-specific structural variation and for interpreting patterns of diversity within and between rice subpopulations. Here, we present a chromosome-scale genome assembly and comprehensive annotation of SM. This resource provides a foundation for comparative genomic analyses, functional genomics, characterization of structural variation, and population-scale investigations.

## Materials and Methods

### Plant material, DNA extraction and Genome sequencing

Nucleus seeds of *Oryza sativa* ssp. *indica* cv. Samba Mahsuri (BPT5204; SM) were obtained from the Agricultural Research Station, Bapatla (Andhra Pradesh, India). Seeds were germinated under controlled conditions, and young leaves from 2-week-old seedlings were used for genomic DNA extraction. Short-read DNA was prepared using the CTAB method (Doyle & Hortorium, 1991). DNA integrity was verified on 0.8% agarose gels, and concentration was measured using a Qubit 3 fluorometer (Invitrogen). Short-read libraries (500 ng DNA) were prepared using the Nextera DNA Flex kit (Illumina Inc., San Diego, CA, USA) following the manufacturer’s instructions, and sequenced on the Illumina NovaSeq 6000 platform in paired-end mode.

For long-read sequencing, approximately 5 µg of sheared genomic DNA (10-25 kb fragments) was used to generate SMRTbell libraries using the SMRTbell Express Template Prep Kit 2.0 (Pacific Biosciences, Menlo Park, CA, USA) and sequenced on the PacBio Sequel II system (Wenger et al., 2019). Ultra-high-molecular-weight (UHMW) DNA was extracted using the Bionano Prep SP Tissue and Plant DNA Isolation Kit, labeled with DLE-1 enzyme, and analyzed on the Bionano Saphyr system according to the manufacturer’s protocols (Bionano Genomics, 2020).

### Read processing and quality control

Details of sequencing platforms, coverage, and library types are provided in **Supplementary Data 1**. Illumina paired-end reads were quality-checked using FASTQC (v0.11.9) and adapter/quality trimmed with Trim Galore! (v0.6.7), applying a Q20 quality threshold and minimum read length of 30 bp. PacBio subreads were processed in SMRT Link (v10.1) to generate high-fidelity (HiFi/CCS) reads using the CCS algorithm with default parameters. Bionano optical mapping data were assembled, filtered, and de-novo consensus genome maps were generated using Bionano Access (v1.7) and the associated Solve pipeline.

### Nuclear Genome assembly

Genome size was estimated using Jellyfish (v2.3.0, k = 21). Short-read *de novo* assemblies were generated using Velvet (v1.2.10) across multiple k-mer values. The CCS reads were assembled using three long-read assemblers, WTDBG2 (v2.2), Hifiasm (v0.16.1), and HiCanu (v2.1.1) with default parameters. For HiCanu, the estimated genome size was set to 400 Mb, while WTDBG2 and Hifiasm were run using their standard presets for HiFi reads. Hybrid assemblies integrating Illumina short reads and HiFi long reads were generated using MaSuRCA (v3.3.4) and HASLR (v0.8), both executed with default settings, supplying an estimated genome size of ∼400 Mb when required. Assembly completeness was evaluated with BUSCO (v3.0; embryophyta odb9) via gVolante.

### Polishing and scaffolding

High-quality Illumina reads were aligned to the Hifiasm assembly using BWA (v0.7.17), and polishing was performed with Pilon (v1.24). Scaffolding was achieved by aligning polished contigs to optical maps using hybridScaffold.pl (Bionano Access). Chromosome-scale pseudomolecules were constructed by ordering super-scaffolds based on alignments to the *O. sativa* Nipponbare and R498 references using minimap2 (v2.24) and D-Genies (v1.4); gaps between adjacent scaffolds were filled with 100 bp Ns.

### Organelle Genome Assembly

Chloroplast genome assembly was performed using GetOrganelle (v1.7.7.1) with the *embplant_pt* seed database. Paired-end reads (mean length ∼140 bp; Phred+33) were filtered, and ∼4.3 million read pairs were retained for assembly. Mapping was done using Bowtie2 (v2.5.4), and assembly was carried out with SPAdes (v4.2.0) using k-mers 21, 55, 85 and 115. The final FASTG graph was disentangled into circular plastome configurations, producing two flip-flop SSC orientations with an average base coverage of ∼495×. Assembly graphs were examined in Bandage (v0.8.1), and the complete plastome sequence was extracted from the FASTG output. D-Genies was employed to compare the assembled sequence with the IRGSP 1.0 and R498 plastome assemblies, and to reorder the sequence, after which the assembly was re-examined in Bandage.

The mitochondrial genome of SM was assembled using three complementary strategies. Illumina paired-end reads were first processed with GetOrganelle (v1.7.7.1), using the plant mitochondrial reference seed database and settings optimized for mitochondrial genomes, including read subsampling to limit excessive depth and dedicated graph-disentangling steps to resolve alternative sub-genomic circles. PacBio CCS reads were then assembled using MitoHiFi (v3.2.3), run in plant mode with a curated *Oryza* mitochondrial reference sequence to guide read recruitment, enforce high-accuracy read matching, and enable integrated mitochondrial gene annotation via MITOS. Finally, a hybrid assembly was generated using Unicycler (v0.5.1) in its high-stringency mode designed to resolve complex graph structures by combining long-read continuity with short-read polishing accuracy.

### Repeat annotation and telomere/centromere identification

Telomeric repeats (AAACCCT)n were identified using tidk (v0.2.1). Centromeric RCS2 repeats (GenBank AF058902) were localized by BLASTN. A *de novo* repeat library was generated with RepeatModeler (v2.0.3), and repeats were annotated and masked using RepeatMasker (v4.1.5).

### Genome Annotation

Total RNA was isolated from root, shoot, leaf, flag leaf, and panicle tissues collected across seedling, tillering, and reproductive stages, and RNA integrity was confirmed using a Bioanalyzer. Pooled RNA was used for PacBio Iso-Seq library preparation and sequencing on the Sequel II platform, while individual tissue samples were processed using the TruSeq Stranded Total RNA with Ribo-Zero Plant kit and sequenced on the Illumina NovaSeq 6000. Iso-Seq (v3.0) subreads were processed to obtain full-length high-quality isoforms, and short-read RNA-seq data were aligned to the SM genome using STAR and assembled with StringTie (v2.2.1). Iso-Seq isoforms and Illumina-based transcripts were merged to generate a non-redundant transcriptome sequence that was subsequently used as evidence for structural gene annotation. Gene prediction incorporated multiple approaches, including evidence-based transcript models, *de novo* predictions from GeneMark, Augustus, and GlimmerHMM and an integrated RNA-seq and protein-supported annotation using Braker3. Completeness of each dataset was assessed using BUSCO (embryophyta odb10), and standard sequence statistics were computed for all predictors.

Functional annotation of predicted protein-coding genes was performed using Panzzer2 and KofamKOALA, and analyses for Augustus and Braker3 gene models were carried out independently. Peptide sequences from each gene prediction set were submitted to Panzzer2, which conducted homology-based searches against integrated reference protein databases and assigned high-confidence functional descriptions and Gene Ontology (GO) terms covering

Molecular Function, Biological Process, and Cellular Component categories. In parallel, the same peptide datasets were annotated with KofamKOALA, which uses profile HMMs and adaptive score thresholds from the KEGG Orthology database to assign KEGG Ortholog (KO) identifiers and pathway annotations.

Organelle genome annotation for the chloroplast was performed using the GeSeq tool within Chlorobox, with the sequence source set as “plastid (land plants)” and minimum BLAT identity thresholds of 25% for proteins and 85% for CDS, tRNA, and rRNA features. The same tool was used with the same thresholds to annotate the mitogenome assembly generated using Unicycler, and sequence source was set as “mitochondrial”.

Non-coding RNA (ncRNA) genes were identified using Infernal (v1.1.5). Structured ncRNAs were detected by searching the assembled SMv1.0 genome against the Rfam covariance model database, applying curated score thresholds to ensure high-confidence matches. This allowed the annotation of major Rfam ncRNA classes, including ribosomal RNAs, small nuclear RNAs, small nucleolar RNAs, and other conserved RNA families. Additionally, tRNA genes were annotated independently using tRNAscan-SE (v2.0.12), which identified canonical tRNAs and recorded their genomic coordinates and classification details.

### Variant discovery and annotation

Whole-genome sequence comparisons were performed by aligning the query genome (IRGSP 1.0/ R498) against the SM genome assembly as a reference using Minimap2, which provided a base for detecting sequence-level and structural variation. For simple variant calling, Single Nucleotide Polymorphisms (SNPs) and small Insertions/Deletions (InDels) were detected using Bcftools (v1.11) mpileup, generating high-confidence variant calls for sequence-level differences between genomes and the distribution of variants was visualized in 1 Mb windows using CMplot (v4.4.1). Structural variants (SVs), including inversions, translocations, and duplications, were identified using SyRI (v1.6.3), enabling assessment of genomic differences in location, orientation, and copy number. Plotsr (v1.1.1) was run, providing comprehensive maps of genomic rearrangements and structural variant distributions across the SM genome. Identified variants, both structural and sequence-level, were annotated using SnpEff (v5.2). Additionally, dot plot alignments between the SM superscaffolds and publicly available *O. sativa* assemblies generated using D-Genies for ordering SM’s pseudomolecules were zoomed in to visualize the IMI (Inversion-Match-Inversion) region.

### Synteny detection, duplication analysis, and evolutionary rate estimation

Collinear genomic regions were identified within the SM genome and between SM and the reference rice genomes Nipponbare (IRGSP-1.0) and R498 using a unified MCScanX-based workflow. Protein sequences derived from curated Braker3 gene models were used for all analyses. Sequences shorter than 30 amino acids or containing internal stop codons were excluded. For each comparison (SM-SM, SM-Nipponbare, and SM-R498), BLASTP similarity searches were performed using BLAST+ with an E-value threshold of 1 × 10 . Gene coordinate files were generated from GFF3 annotations, and BLASTP outputs together with gene positions were supplied to MCScanX (v1.0.0) to identify collinear blocks. Only blocks supported by multiple, consecutively ordered anchor gene pairs were retained, thereby excluding isolated or spurious matches. In the SM-SM analysis, anchor pairs represent duplicated paralogous genes, whereas in SM-Nipponbare and SM-R498 comparisons, anchor pairs represent putative orthologs defining conserved synteny. For all SM-SM collinear blocks, anchor gene pairs were extracted using custom Python scripts. Coding sequences corresponding to each gene were retrieved directly from the SM genome assembly using gffread and aligned codon-wise. Synonymous (Ks) and nonsynonymous (Ka) substitution rates, as well as Ka/Ks ratios, were estimated using KaKs_Calculator 2.0 under the Yang-Nielsen model. Gene pairs with unreliable or saturated estimates were excluded from downstream analyses. Genome-wide distributions of Ka, Ks, and Ka/Ks values were summarized using histograms and violin plots. Anchor gene pairs were associated with their corresponding collinear blocks, and block-level summary statistics, including the number of contributing gene pairs and mean Ks values, were calculated. Duplication ages were inferred from Ks values assuming a synonymous substitution rate of 6.5 × 10 substitutions per site per year, estimated from coding regions of adh1 and adh2 genes in grasses (Gaut et al., 1996; Ma & Bennetzen, 2004). Gene pairs with Ka/Ks > 1 were identified as candidates for accelerated evolution. Functional annotations were integrated from Panzzer2 and KofamKOALA outputs. Genome-wide and chromosome-level synteny patterns within SM and between SM, Nipponbare, and R498 were visualized using MCScanX plotting utilities and the SynVisio platform. Gene position files and anchor gene pair lists were uploaded to SynVisio to generate linear and ribbon-style synteny plots.

### Population genomic analyses

Whole-genome short-read sequencing data for 533 cultivated rice accessions were obtained from the RiceVarMap2 dataset via the NCBI SRA. Raw reads were independently mapped to the SM reference genome using standard alignment workflows. Joint variant calling across all accessions was performed using bcftools mpileup followed by bcftools call (bcftools v1.11; htslib v1.13). The resulting VCF was compressed and indexed using tabix, and variant-level summary statistics including allele counts, allele frequencies, minor allele frequency (MAF), missingness, and sample number were annotated using the bcftools +fill-tags plugin. SNPs were filtered to retain high-confidence variants with missingness ≤ 10% and MAF ≥ 0.05. Sample identifiers were standardised using bcftools reheader. Filtered SNPs were converted to PLINK binary format using PLINK (v1.9). Linkage disequilibrium pruning was performed using a sliding window approach (200 SNP window, 50 SNP step size, r² threshold 0.2) to generate a set of approximately independent markers. Population structure was assessed using principal component analysis (PCA) implemented in PLINK (v1.9). Eigenvectors generated by PLINK were used for downstream visualisation using custom Python scripts. Exploratory analyses were performed using both pruned and unpruned datasets; however, all PCA results presented in this study are based on LD-pruned SNPs to minimise the influence of linked variation. Phylogenetic relationships among accessions were inferred by computing pairwise identity-by-state (IBS) distances using PLINK (v1.9) on an LD-pruned, genome-wide SNP dataset. To assess locus-specific phylogenetic relationships, IBS distance matrices and phylogenetic trees were additionally constructed using SNPs restricted to the chromosome 6 IMI interval, following the same LD-pruning and distance estimation procedures. Neighbor-Joining trees were constructed in R from the distance matrices and exported in Newick format for visualisation using iTOL. To investigate population genetic patterns associated with the chromosome 6 IMI locus, nucleotide diversity (π), Tajima’s D, and genetic differentiation (FST) were quantified across three genomic intervals on chromosome 6: the IMI region, an upstream (pre-IMI) region, and a downstream (post-IMI) region for the two indica clusters obtained in the IMI-region PCA plot. All analyses were based on SNPs from the joint-genotyped dataset. Accessions were assigned to two genetic clusters based on IMI-specific population structure. Cluster-specific VCF files were generated using bcftools (v1.11), and high-confidence biallelic SNPs were retained using the following criteria: SNPs only, biallelic sites, QUAL > 30, and total read depth > 10. Nucleotide diversity (π) was estimated separately for each cluster using VCFtools (v0.1.17) in non-overlapping 50 kb windows. Tajima’s D was calculated using the same window size, and windows with fewer than 20 segregating sites or undefined values were excluded. Genetic differentiation between the two clusters was quantified using Weir and Cockerham’s FST as implemented in VCFtools (v0.1.17), and was calculated both at individual SNPs and in non-overlapping 50 kb windows. Windows with missing or undefined FST values were excluded prior to visualisation. For each genomic interval, window-based estimates of π, Tajima’s D, and FST were merged based on genomic coordinates. Minor coordinate offsets introduced by window-based calculations were corrected prior to merging. Final summary datasets were visualised using custom R scripts.

## Results

### Genome characteristics of Samba Mahsuri

K-mer-based analysis using GenomeScope (v1.0; *k* = 21) estimated a haploid genome size of ∼338 Mb, with low heterozygosity (0.045–0.058%) and approximately 26% repetitive content (***Figure S1***). The model indicated a high proportion of unique sequence (∼73.6%) and a low sequencing error rate (∼0.74%). The k-mer–based estimate is lower than the final assembly size, consistent with the known tendency of k-mer models to underestimate genome size in repeat-rich plant genomes.

### Nuclear Genome Assembly

Initial short-read assemblies of the SM genome generated using Velvet were highly fragmented, with total assembly sizes reaching up to 354 Mb and a maximum N50 of 11.4 kb (***Supplementary Data 1***). Despite this fragmentation, the largest short-read assembly recovered 96.8% of BUSCO genes, indicating that most gene content was captured. However, the limited contiguity of these assemblies precluded their use as a reference genome. Hybrid assemblies using the PacBio CCS and Illumina reads were generated by HASLR and MaSuRCA in an attempt to improve contiguity. While the HASLR assembly remained fragmented, comprising 6,602 contigs with a total length of 273.8 Mb and an N50 of 75 kb, MaSuRCA produced a substantially more contiguous assembly consisting of 768 contigs (N50 = 1.15 Mb; total length = 396.7 Mb) with high completeness (BUSCO = 97.8%), although gaps were present in 31 contigs. Long-read assemblies, using only the PacBio CCS reads (∼139× raw coverage; mean read length ∼14 kb) with Wtdbg2, HiCanu, and Hifiasm, resulted in Hifiasm giving the most contiguous assembly. The Wtdbg2 assembly was highly fragmented (4,058 contigs; N50 = 191 kb; BUSCO = 96.6%), while HiCanu improved contiguity (1,370 contigs; N50 = 674 kb; BUSCO = 97.0%). Hifiasm assembly, comprised of 767 contigs with an N50 of 1.29 Mb and a total assembly length of 401.0 Mb (BUSCO = 97.3%). This assembly included 600 primary contigs and 167 alternate haplotype-phased contigs. Given its superior contiguity and completeness, the Hifiasm assembly was selected for polishing. Alignment of Illumina short reads followed by correction with Pilon resolved 37,860 base-level errors and improved BUSCO completeness to 97.9%. The polished Hifiasm assembly (400.94 Mb; N50 = 1.29 Mb) was retained for downstream analyses (***Figure S2***).

### Optical map-guided scaffolding and pseudomolecule construction

In order to get closer to a chromosomal level assembly, Bionano optical mapping using the DLE-1 enzyme was carried out, which generated high-quality molecules at approximately 345× coverage, with a mean molecule length of 214 kb and an average label density of 13 sites per 100 kb. Assembly of these molecules produced 63 consensus genome maps totaling 404 Mb, with a mean map length of 6.4 Mb. Aligning the polished Hifiasm contigs to the optical maps enabled anchoring of 449 contigs into 30 super-scaffolds. The scaffolded assembly spanned 7.98 Mb, with an N50 of 16.3 Mb and an L50 of 10. A total of 831 gaps, accounting for 3.7 Mb of Ns, remained between contigs. BUSCO analysis indicated high completeness (C: 97.7% [S: 96.9%, D: 0.8%]), reflecting minimal loss of gene content during scaffolding. To generate chromosome-scale pseudomolecules, the super-scaffolds were ordered and oriented using synteny with the Nipponbare, R498 and other reference genomes in D-genies (***Figure 1***, ***S3***). Adjacent super-scaffolds were connected by inserting blocks of 100 Ns to represent unresolved gaps. The final pseudomolecule assembly comprised 12 chromosomes spanning 394.99 Mb, with an N50 of 31.7 Mb (***Table S1, S2***).

**Figure 1.**
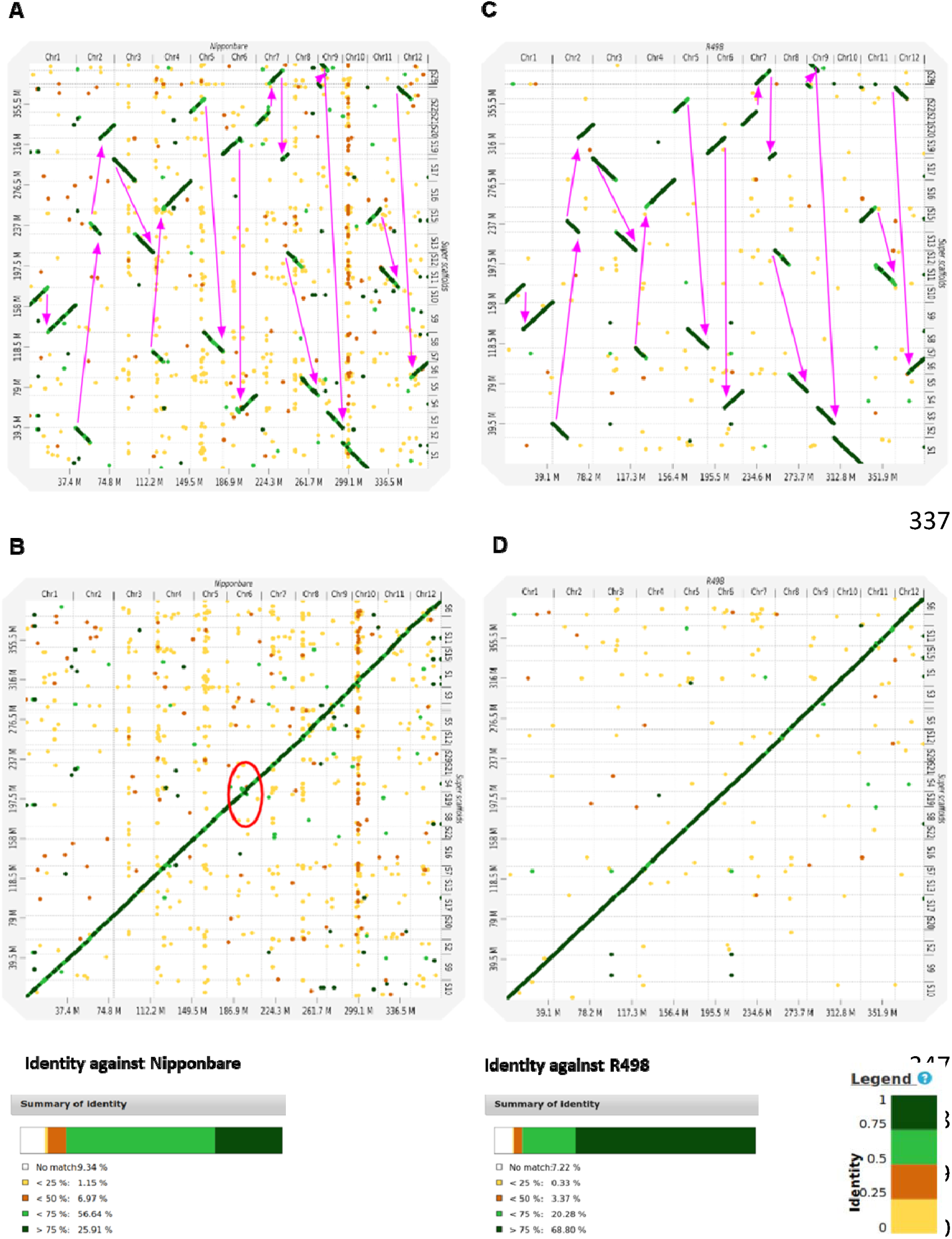
Dot plot comparison of SM super-scaffolds with reference Oryza sativa genomes. ***(A)*** Alignment of SM super-scaffolds against the Nipponbare genome (IRGSP-1.0) showing overall one-to-one correspondence with dispersed low-identity signals. (***B***) The same alignment with SM super-scaffolds reordered according to Nipponbare chromosomal coordinates, revealing a largely conserved diagonal pattern and a clear inversion signal on chromosome 6 (circled). (***C***) Alignment of SM super-scaffolds against the R498 genome, showing similar large-scale collinearity with expected variation in local identity. (***D***) Reordered SM super-scaffolds according to R498 chromosome order, improving visualization of collinear relationships. In all panels, coloured lines and dots indicate pairwise identity (green: high; yellow/orange: lower), and diagonal tracks denote conserved synteny.

### Chromosome-scale structure of SM pseudomolecules

Chromosome-wise comparison revealed that the SM assembly (394.9 Mb) is larger than both the Nipponbare (373.2 Mb) and R498 (390.3 Mb) genomes (***Figure S4, Table S3***). Overall chromosome structure and ordering were highly conserved across the three genomes. However, modest chromosome-specific expansions were observed in SM, particularly on Chr4, Chr5, Chr7, Chr8, Chr9, and Chr 11. In contrast, R498 exhibited the longest chromosomes for Chr1, Chr2, Chr3, Chr6, and Chr10, while Nipponbare retained a slightly longer Chr12. Despite these differences, the relative distribution of chromosome sizes across the three genomes remained highly consistent, supporting the accuracy and structural integrity of the SM pseudomolecule reconstruction.

### Organelle genome assembly: plastome and mitogenome reconstruction

The plastid genome was assembled using GetOrganelle, yielding a complete circular plastome with the canonical quadripartite structure comprising the large single-copy (LSC), small single-copy (SSC), and inverted repeat (IR) regions (***Figure S5A***). All junctions were resolved unambiguously, with no detectable structural inconsistencies. In contrast, the mitochondrial genome could not be resolved into a single contiguous structure despite the use of multiple assembly approaches, including GetOrganelle, MitoHifi, and Unicycler. All methods recovered complex, multi-isoform assemblies characteristic of highly recombinogenic plant mitogenomes, and no single configuration could be designated as a canonical mitochondrial chromosome. The alternative graph structures and isoforms generated by each assembler are presented in ***(Figure S5B-D)*.**

### Repeat repertoire and structural features of the SM genome

The SM pseudomolecules were partitioned into 10 kb windows, and the plant telomeric motif AAACCCT was identified using tidk. All chromosomes showed enrichment of telomeric repeats toward at least one end, and chromosomes 1, 2, 3, 4, 5, 6, and 11 displayed tandem AAACCCT arrays at both ends (***Figure S6***), indicating that terminal chromosomal regions are well represented in the assembly. Centromeric regions were localized using RCS2 repeats, which are conserved centromeric elements in *Oryza*. Clusters of RCS2 matches delineated putative centromeres across all SM pseudomolecules (***Figure S6, Table S4***). Isolated RCS2 hits outside the main centromeric clusters were detected on chromosomes 3 and 8, while chromosome 9 contained multiple gapped RCS2 arrays, likely reflecting the structural complexity and limited resolvability of highly repetitive centromeric regions. In chromosomes assembled from multiple super-scaffolds, RCS2-based centromeric clusters were consistently confined to a single region near the end of the first super-scaffold, a pattern consistent with expected centromere localization given the limited resolvability of highly repetitive centromeric sequences by both contig assembly and DLE-1–based optical mapping. Repeat annotation using RepeatModeler indicated that 50.9% of the SM genome is composed of repetitive sequences (***Table S5***). Interspersed repeats accounted for 49.24% of the genome, dominated by retroelements (24.26%), primarily long terminal repeat (LTR) elements (23.2%). Among these, Gypsy/DIRS1 elements were the most abundant (17.21%), followed by Ty1/Copia elements (2.88%). DNA transposons contributed 3.7% of the genome, while unclassified repeats comprised 21.28%. Additional repeat categories included simple repeats (1.11%), low-complexity sequences (0.13%), satellites (0.05%), and small RNAs (0.11%). All identified repeats were masked prior to gene prediction. Repeat density was enriched in pericentromeric regions, consistent with patterns observed in other indica genomes (***Figure 2***).

**Figure 2.**
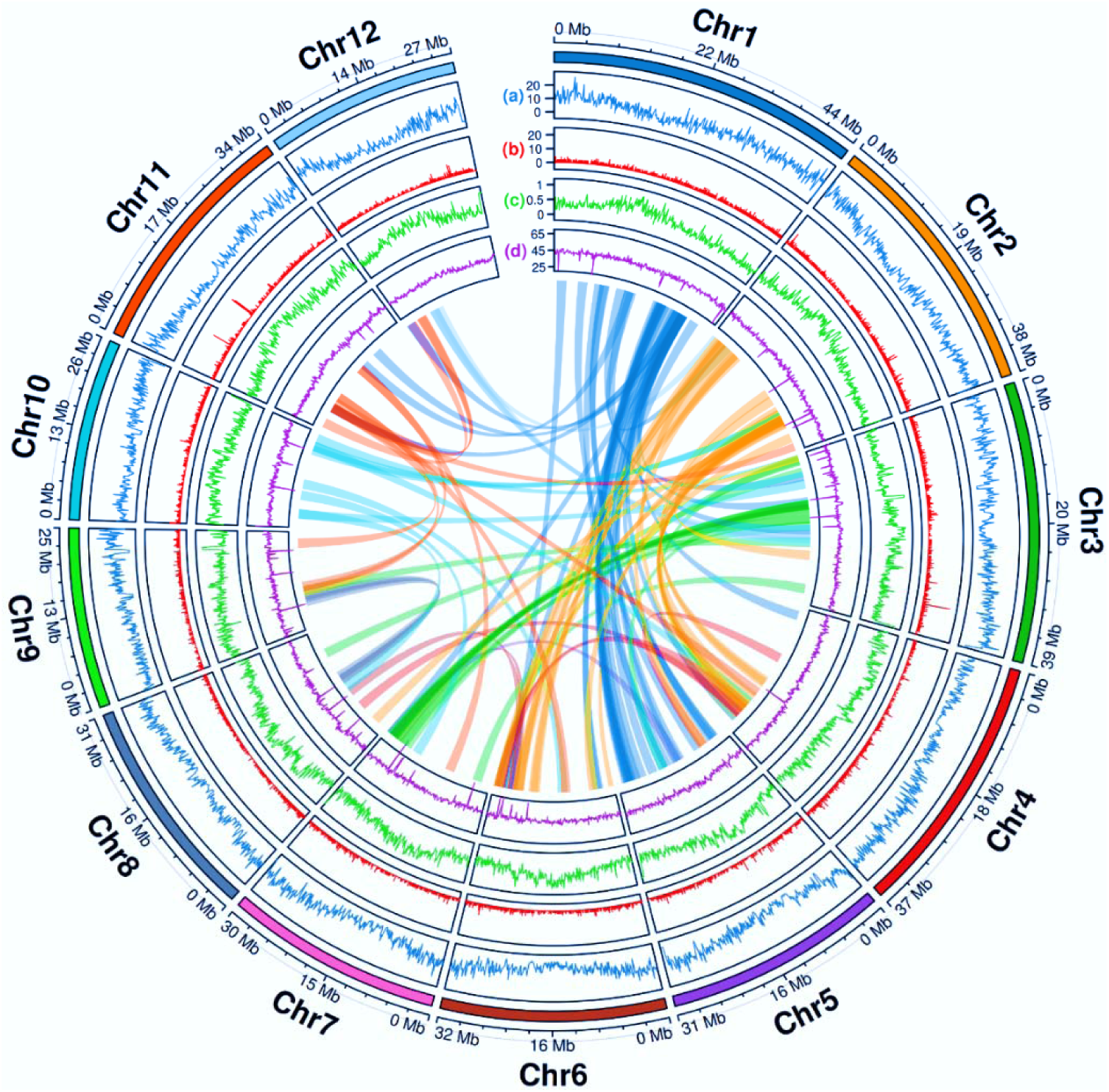
A Circos plot representation of the SM genome. Outer tracks show ***(a)*** protein-coding gene density (100 kb windows), ***(b)*** non-coding RNA density (10 kb), ***(c)*** repeat density (100 kb), and ***(d)*** GC content (100 kb). Inner ribbons depict intragenomic syntenic relationships, highlighting conserved collinear blocks across chromosomes and providing a framework for the segmental duplication analyses presented in subsequent sections.

### Protein-coding gene models in SM

Long-read transcriptome sequencing using IsoSeq generated 36,759 high-quality full-length transcripts supported by at least two CCS reads. Genome alignment and isoform collapsing yielded a non-redundant set of 14,667 unique IsoSeq isoforms. Transcript assemblies from Illumina short-read RNA-seq data contributed additional isoforms from individual samples (**Supplementary Data 1**). Merging IsoSeq- and Illumina-derived transcript sets produced a combined transcriptome comprising 35,686 isoforms originating from 24,939 genes. These transcript assemblies were used as supporting evidence for genome annotation. Genome-wide gene prediction was performed independently using ab initio and evidence-guided approaches (***Table S6***). BUSCO analysis indicated that transcript-derived models alone achieved 85.3% completeness. Among ab initio predictors, Augustus and GlimmerHMM recovered 93.0% and 89.3% complete BUSCOs, respectively, while GeneMark retrieved substantially fewer complete BUSCOs (10.0%). Based on its higher completeness and well-balanced gene model structure, the Augustus prediction set was retained as the final ab initio gene dataset for downstream analyses. Evidence-guided gene prediction was performed using Braker3, which integrates RNA-seq alignment evidence with ab initio prediction to infer gene models across the entire genome, including regions not represented in transcript assemblies. Braker3 exhibited the highest overall completeness, identifying 97.4% complete BUSCOs with only 2.35% missing, and predicted 31,138 genes corresponding to 35,069 transcripts. Accordingly, the Braker3 output was retained as the final gene model set. Overall, integration of long-read and short-read transcript evidence with complementary ab initio and evidence-guided gene prediction approaches yielded a high-confidence and well-supported annotation of the SM nuclear genome.

Functional annotation was performed independently on the final ab initio and evidence-based nuclear gene model sets. The Augustus gene set comprised 59,152 predicted gene models, of which 31,923 were assigned functional descriptions based on homology and annotation evidence. Gene Ontology (GO) terms were distributed across all three major categories, with 22,598 genes in Molecular Function, 18,086 in Biological Process, and 15,433 in Cellular Component categories. The evidence-based Braker3 annotation predicted 31,138 genes corresponding to 35,069 transcripts. Functional annotation using Panzzer2 assigned functional descriptions to 26,250 Braker3 gene models. GO annotation of the Braker3 gene set revealed 20,778 genes associated with Molecular Function terms, 19,029 with Biological Process terms, and 21,871 with Cellular Component terms, reflecting broad functional coverage across core cellular and metabolic processes. Parallel annotation using KofamKOALA found KEGG orthologs to a substantial fraction of predicted gene models from both datasets, identifying them in conserved biochemical and regulatory pathways. Core pathways, including carbohydrate metabolism, amino acid biosynthesis, energy production, and lipid metabolism, were well represented. Together, the Panzzer2 and KofamKOALA annotations provide consistent functional support for both ab initio and evidence-based gene models and enable downstream pathway-level and comparative genomic analyses. Protein-coding genes were unevenly distributed across chromosomes, with higher densities toward distal chromosomal regions (***Figure 2***).

### Non-coding RNA encoding Genes in the Samba Mahsuri genome

There were 8,179 ncRNA loci distributed across all 12 chromosomes of the SM genome (***Supplementary Data 1***). The annotated set includes tRNAs, ribosomal RNAs, microRNAs, and other conserved structured RNAs, indicating comprehensive recovery of major ncRNA classes. A total of 606 tRNA genes, including two selenocysteine tRNAs, were distributed across chromosomes, consistent with expectations for a complete rice nuclear genome. Ribosomal RNA loci, on the other hand, exhibited strong chromosomal clustering. In particular,5S rDNA repeats were localized exclusively to chromosome 11, forming multiple tandem clusters within the centromeric or secondary constriction region. This clustering of 5s rDNA on chromosome 11 was confirmed by comparing with Nipponbare and R498. Analysis of 5S rDNA repeat structure revealed genome-specific differences in intergenic spacer length. The average spacer length in SM (∼462 bp) was longer than in Nipponbare (∼220 bp) and comparable to R498 (∼449 bp), indicating divergence in repeat unit organization among rice lineages. The microRNA repertoire showed marked family-specific expansion. The MIR812 family accounted for the majority of annotated miRNA loci, while several other miRNA families were present at moderate copy numbers. Many miRNA families occurred as single or low-copy loci. In addition, numerous snoRNA and snRNA families were recovered, most at low copy number, consistent with their conserved cellular roles. Intragenomic synteny revealed extensive collinearity between homologous chromosomal segments (***Figure 2***).

### Organelle genome annotation

The SM plastid genome (∼134.5 kb) was annotated using GeSeq, yielding 85 unique protein- coding genes, 24 tRNA genes, and the four canonical ribosomal RNA genes (*16S, 23S, 4.5S,* and *5S rRNAs*). Gene order and content were highly conserved relative to other *Oryza* chloroplast genomes, and the plastome exhibited the typical quadripartite architecture comprising the large single-copy (LSC), small single-copy (SSC), and inverted repeat (IR) regions (***Figure S5***).

Annotation of mitochondrial contigs generated by the Unicycler hybrid assembly identified the expected complement of rice mitochondrial genes. These included core components of the respiratory chain, such as *NADH dehydrogenase* (*nad* genes), *cytochrome c oxidase* (*cox* genes), *ATP synthase* (*atp* genes), and *cytochrome b* (*cob*), along with the canonical set of mitochondrial tRNAs and several lineage-typical hypothetical ORFs. Fragmented ORFs and duplicated features were also detected, consistent with the multipartite and recombination-driven organization characteristic of angiosperm mitogenomes.

Comparison of annotated SM mitochondrial genes with the Nipponbare mitogenome revealed broad conservation of the core gene set. Shared genes included *nad1*, *cox3*, *orf25*, *orf152a*, and multiple tRNAs (e.g., *trnP*, *trnM*, *trnH*, and *trnW*). However, the SM assembly contained ORF fragments and tRNA variants not present in Nipponbare, whereas Nipponbare retained genes or ORFs absent in SM, including *rpl16*, *rps11*, *orf241*, and *nad4L*. Despite these differences, the conserved complement of core mitochondrial genes confirms that the SM mitogenome retains all essential mitochondrial gene functions characteristic of cultivated rice.

### Syntenic duplication history and selective constraint in the SM genome

To characterize the evolutionary history and selective pressures acting on duplicated genes in the SM genome, we analyzed 586 high-confidence anchor paralog pairs identified from intragenomic collinearity detected by MCScanX (***Figure 3***). These paralogs represent conserved syntenic duplicates with reliable alignments and substitution rate estimates. Genome-wide distributions of nonsynonymous (Ka), synonymous (Ks), and Ka/Ks ratios revealed uniformly low Ka values and strongly right-skewed Ka/Ks ratios, with median Ka = 0.145, Ks = 1.00, and Ka/Ks = 0.126, indicating pervasive purifying selection acting on retained duplicates. The majority of paralogs showed Ka/Ks < 0.3, while only five gene pairs exhibited Ka/Ks > 1, suggesting limited instances of potential functional divergence (***Table S7***).

**Figure 3.**
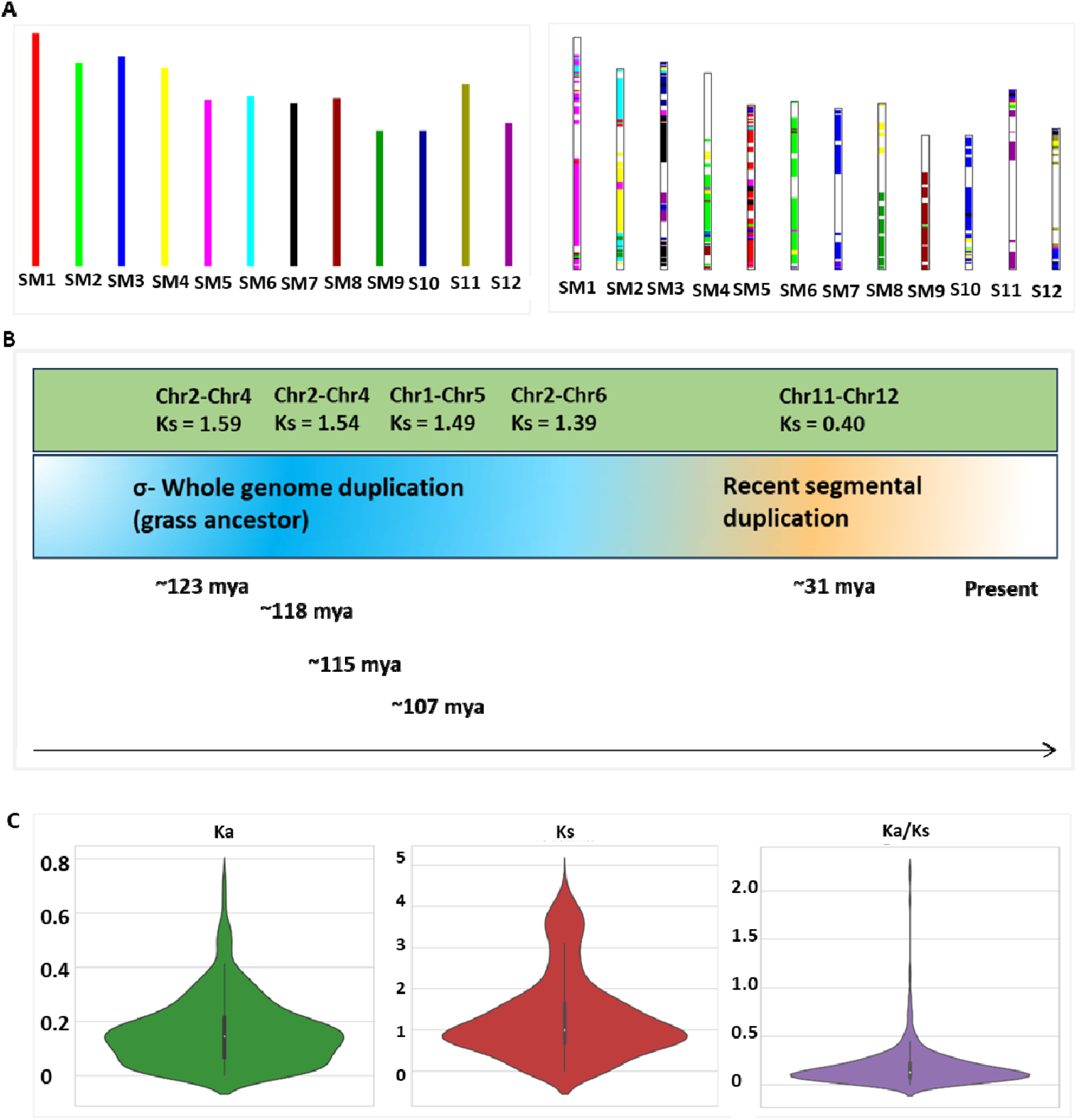
Intragenomic duplication landscape and evolutionary rates in the SM genome. (***A***) Left: ideograms of the 12 SM chromosomes. Right: intragenomic syntenic segments detected by MCScanX, illustrating dispersed and fragmented paleoduplication remnants rather than recent large-scale duplication. (***B***) Summary of major duplicated chromosomal blocks in the SM genome plotted by mean synonymous substitution rate (Ks) highlighting blocks associated with the ancient σ whole-genome duplication shared across grasses (Ks ≈ 1.4-1.6) and a younger segmental duplication between chromosomes 11 and 12 (Ks ≈ 0.4). (***C***) Distributions of Ka, Ks, and Ka/Ks values for 586 duplicate gene pairs, showing uniformly low Ka, broad Ks variation, and predominantly Ka/Ks < 1, indicating strong purifying selection acting on ancient retained duplicates.

Block-wise aggregation of Ks values grouped these paralogs into six major collinear blocks spanning specific chromosome pairs (***Table S8***). Five blocks, with mean Ks values ranging from 1.39 to 1.59, correspond to ancient paralogy relationships derived from the σ whole-genome duplication shared across grasses (∼100-130 MYA). In contrast, a single block linking chromosomes 11 and 12 displayed a markedly lower mean Ks (∼0.40; ∼30 MYA), consistent with a more recent, lineage-specific segmental duplication within *Oryza*. No evidence was detected for recent whole-genome duplication events in SM. Together, these patterns indicate that the SM genome is dominated by deeply conserved paleoduplications under long-term purifying selection, with limited contribution from younger segmental duplications.

### Genome-wide divergence and structural variation relative to rice reference genomes

Whole-genome comparisons of the SM assembly with the *japonica* reference, Nipponbare and the *indica* reference, R498 revealed asymmetric but structured patterns of sequence and structural divergence. SM harboured approximately 2.39 million variants relative to Nipponbare and ∼0.98 million relative to R498, reflecting the deeper divergence between *indica* and *japonica* lineages. In both comparisons, SNPs constituted more than 99% of variants, with similar transition/transversion ratios (∼2.6) and comparable missense-to-silent ratios (∼1.15), indicating broadly similar mutational biases and selective constraints despite differences in overall divergence. Genome-wide SNP density profiles showed heterogeneous divergence along chromosomes in both comparisons, with consistently higher SNP densities observed in the SM-Nipponbare contrast relative to SM-R498 (***Figure 4A-B***). Divergence was non-uniform present as broad, megabase-scale regions rather than as isolated peaks. Whole-genome structural alignments revealed extensive conserved macrosynteny between SM and both reference genomes, accompanied by non-random structural variation (***Figure 4C-D***). Nipponbare showed a higher frequency of inversions, translocations, and non-collinear segments compared to R498, consistent with SM’s closer relationship to the *indica* lineage. Notably, several SM-specific rearrangements were present in R498, indicating structural differentiation not captured by either reference genome.

**Figure 4.**
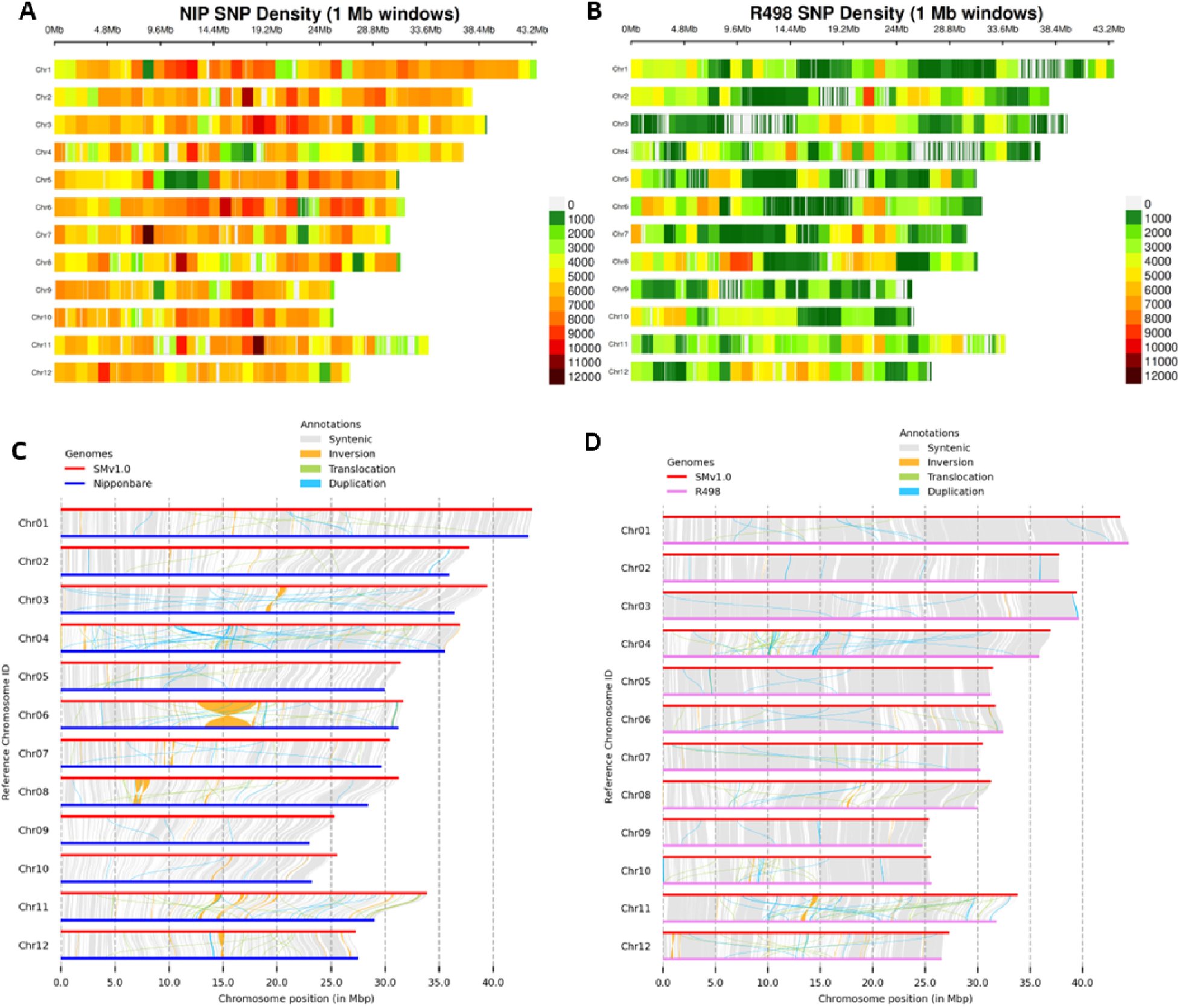
Genome-wide SNP density and structural variation between SM and reference rice genomes. ***(A-B)*** SNP density profiles (1 Mb bins) of the SM assembly relative to *O. sativa* cv. Nipponbare ***(A)*** and *O. sativa* cv. R498 ***(B)*** Warmer colours indicate higher SNP density per bin, highlighting lineage-specific divergence, regions of accelerated evolution, and intervals with reduced variation. Whole-genome structural comparison of SM with ***(C)*** Nipponbare and ***(D)*** R498 using synteny alignments. Syntenic blocks (grey), inversions (yellow), translocations (orange), and duplicated segments (cyan) are shown along SM chromosomes. These comparisons reveal extensive shared macrosynteny across genomes, together with clear cultivar-specific rearrangements, most notably on chromosomes 4 and 11, highlighting structural diversification within *O. sativa*.

Detailed structural variant profiling highlighted several key features of genome-wide divergence. These included (i) large non-aligned segments in SM-Nipponbare (>168 Mb) and SM-R498 (>83 Mb), consistent with the retention of distinct haplotype blocks in SM; (ii) extensive query-side duplications in SM (6,128 relative to Nipponbare and 4,723 relative to R498), encompassing numerous duplicated genomic segments that include genes associated with receptor-like kinases, metabolic functions, and putative resistance pathways; and (iii) highly diverged regions that were approximately threefold larger in the SM-Nipponbare comparison (∼25 Mb) than in SM-R498 (∼8 Mb). (iv) cultivar-specific rearrangements, were noted on chromosomes 4 and 11.

Together, these analyses indicate that while SM is most closely related to the *indica* reference R498, it harbours unique haplotypes and structural configurations that distinguish it from both reference genomes.

### The IMI Block on Chromosome 6

Genome-wide dot plot comparisons revealed extensive macrosynteny between the SM genome and both rice reference genomes, punctuated by localized structural differences (***Figure 5A-B***). One of the most prominent deviations from collinearity was detected on chromosome 6 in the - Nipponbare, whereas the corresponding region showed uninterrupted collinearity in R498. Higher-resolution inspection of chromosome 6 revealed that this region comprises a complex Inversion-Match-Inversion (IMI) configuration in Nipponbare relative to SM, rather than a simple inversion (***Figure 5D***). The IMI structure spans Chr6:12.67-18.53 Mb in SM and corresponds to Chr6:13.12-17.63 Mb in Nipponbare. In contrast, the *indica* reference genome R498 displays a direct collinear orientation across the same interval. Genome-wide synteny visualization confirmed that this structural configuration is restricted to a discrete interval on chromosome 6 and does not extend to other chromosomes (***Figure 5C***). Based on gene annotation, the IMI interval contains 710 predicted genes according to the Augustus ab initio models, whereas the evidence-guided Braker3 annotation identifies 218 genes within the same region. Functional annotations associated with genes located within the IMI interval span diverse biological categories, including DNA-associated processes, stress-related functions, and flowering-related pathways, indicating that the locus encompasses a gene-rich and functionally heterogeneous genomic segment that may be sensitive to structural variation.

**Figure 5.**
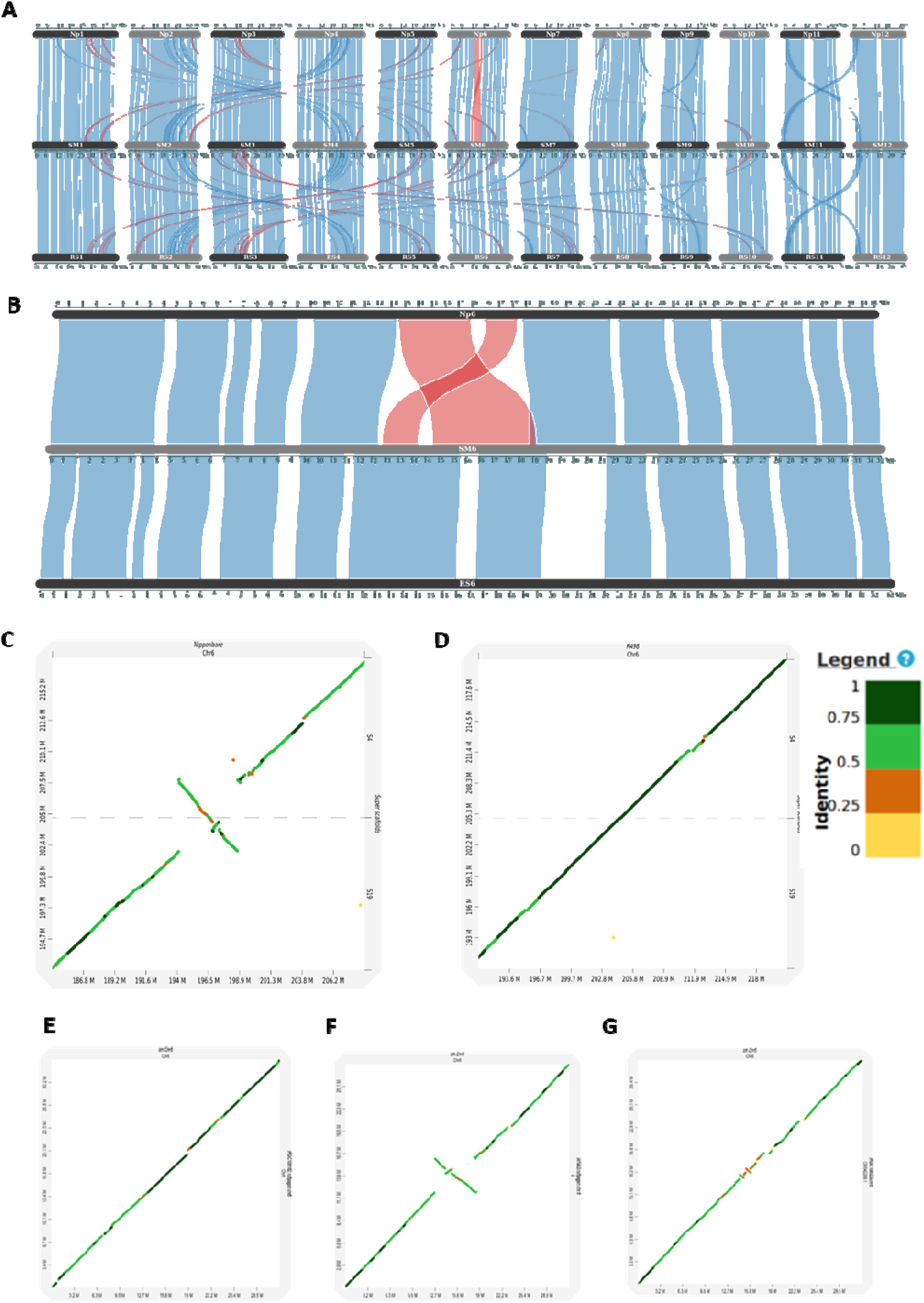
Genome-wide context and locus-specific architecture of the chromosome 6 IMI region. (***A***) Genome-wide synteny comparison between the SM genome (middle track) and the reference genomes Nipponbare (top) and R498 (bottom). Conserved collinear blocks are shown in blue, while structural rearrangements, including inversions, translocations, and duplications, are highlighted in red. ***(B)*** Chromosome-scale synteny view of chromosome 6, showing red. a prominent structural rearrangement in SM relative to Nipponbare centered on the pericentromeric region. ***(C–D)*** Dot-plot alignments of the chromosome 6 IMI interval between SM and Nipponbare (C) and between SM and R498 ***(D)*** SM-Nipponbare comparisons reveal a characteristic inversion-match-inversion (IMI) alignment pattern, whereas SM-R498 shows direct collinearity across the same interval. ***(E–G)*** Dot-plot comparisons of the SM chromosome 6 IMI region with representative wild Oryza genomes, illustrating variation in local alignment patterns, including direct orientation and inversion-associated configurations. Alignment color intensity reflects sequence identity.

Dot-plot comparisons of the chromosome 6 IMI region revealed pronounced structural polymorphisms among cultivated rice and wild relatives. In R498, the locus displayed a direct collinear orientation, whereas the *japonica* reference Nipponbare exhibited a clear inversion-match-inversion (IMI) configuration across the corresponding interval. Comparisons with multiple *Oryza rufipogon* isolates demonstrated that both structural configurations segregate within wild populations. Isolate A showed a direct orientation matching with SM, while isolate B recapitulated the Nipponbare-like IMI structure. Isolate C displayed a more complex pattern, characterized by short inverted segments relative to SM but a complete IMI configuration when aligned to Nipponbare, consistent with nested rearrangements within the larger IMI framework. Together, these comparisons show that both direct and IMI configurations are present among cultivated rice and *O. rufipogon* isolates, with isolate-specific differences in orientation and internal rearrangement patterns at the chromosome 6 locus.

### IMI-specific population structure across cultivated and wild rice accessions

Principal component analysis (PCA) was performed using joint genotyping data from 533 cultivated rice accessions spanning all major *O. sativa* subpopulations, together with four wild accessions (*O. rufipogon* isolates A, B, and C, and *O. nivara*), using the SM genome as the reference (***Figure 6***). IMI-region PCA retained the broad population structure observed at the chromosome scale, but with markedly tighter clustering, separating major rice subpopulations along the primary principal components and distinctly clustering representative *indica*, *japonica*, *aus*, *aro*, and intermediate cultivars, while further resolving discrete *indica* subclusters not observed in whole-chromosome or genome-wide analyses. The wild accessions occupied distinct positions in the IMI PCA. *O. rufipogon* isolate A grouped within the *indica* cluster, whereas isolate B clustered closer to *japonica* accessions. Isolate C formed a separate cluster intermediate between cultivated groups. *O. nivara* occupied a distinct position adjacent to, but clearly separated from, *indica* accessions. In contrast, PCA performed on multiple pre-IMI and post-IMI windows of comparable size, as well as on entire chromosome 6 and chromosome 1, showed substantially weaker or diffuse clustering (***Figure 6B-G***). In these analyses, *indica* substructure was largely unresolved, and separation between cultivated groups was reduced relative to the IMI locus.

**Figure 6.**
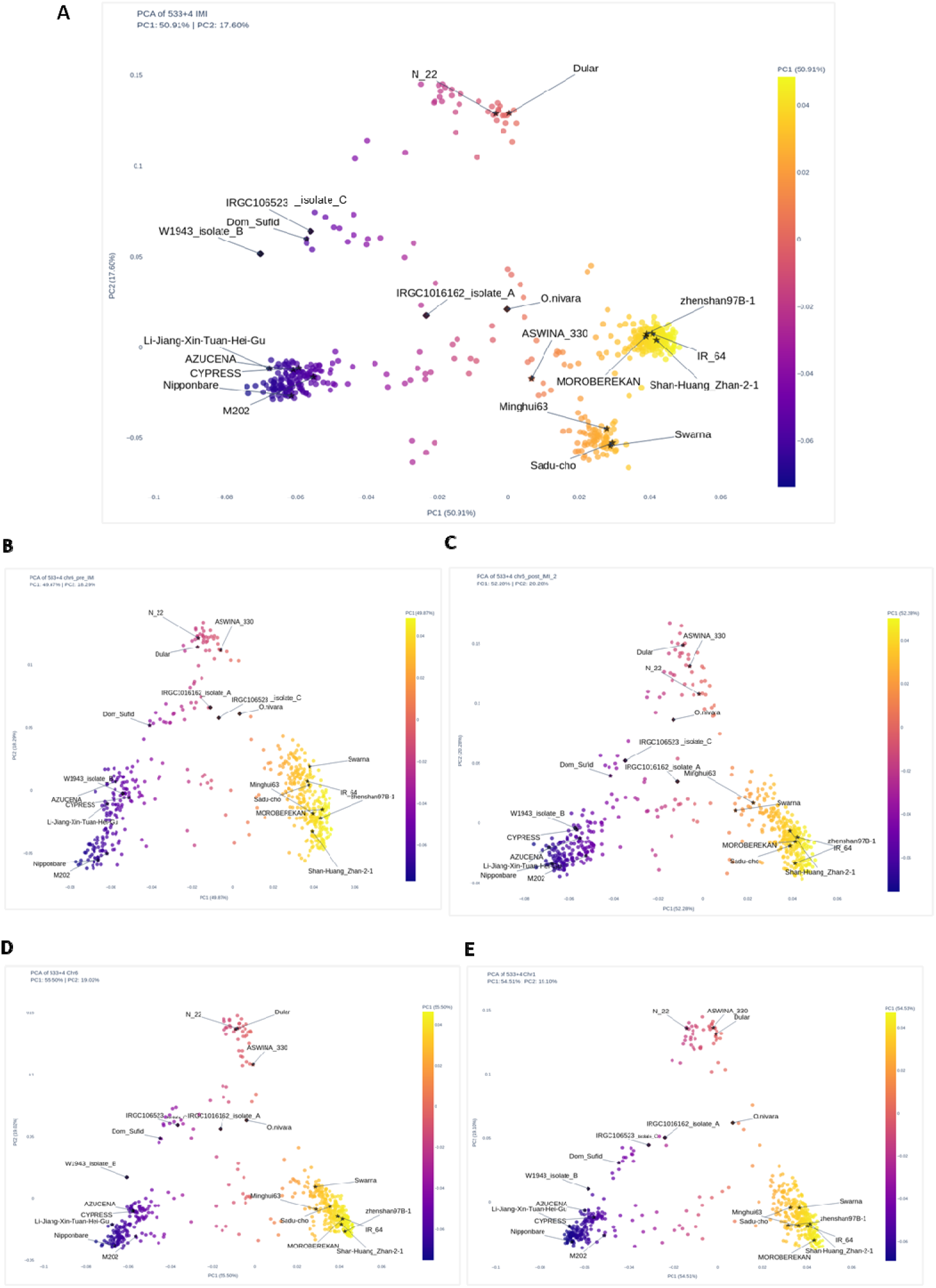
IMI-specific population structure revealed by PCA analysis. Principal component analysis (PCA) of 533 rice cultivars based on joint SM genome as reference. (***A***) (larger) shows PCA performed using genotyping using the g variants from the chromosome 6 IMI region, revealing pronounced population structure, including subdivision within *indica*. (***B, C***) show PCA results for multiple pre-IMI and post-IMI windows of comparable size, while (***F, G***) show PCA for whole chromosome 6 and chromosome 1, respectively. In contrast to the IMI region, flanking and whole-chromosome level analyses show weaker or diffuse clustering, indicating that population structure at the IMI locus is distinct from broader genomic patterns. Cultivars are colored by major rice groups, and a subset of representative accessions from the 533 cultivars is labeled for reference.

Neighbor-joining trees were constructed using genome-wide SNPs and SNPs restricted to the chromosome 6 IMI region (***Figure 7***). The genome-wide tree grouped accessions according to major rice subpopulations, including *indica*, *japonica*, aus, aromatic, and intermediate groups.

**Figure 7.**
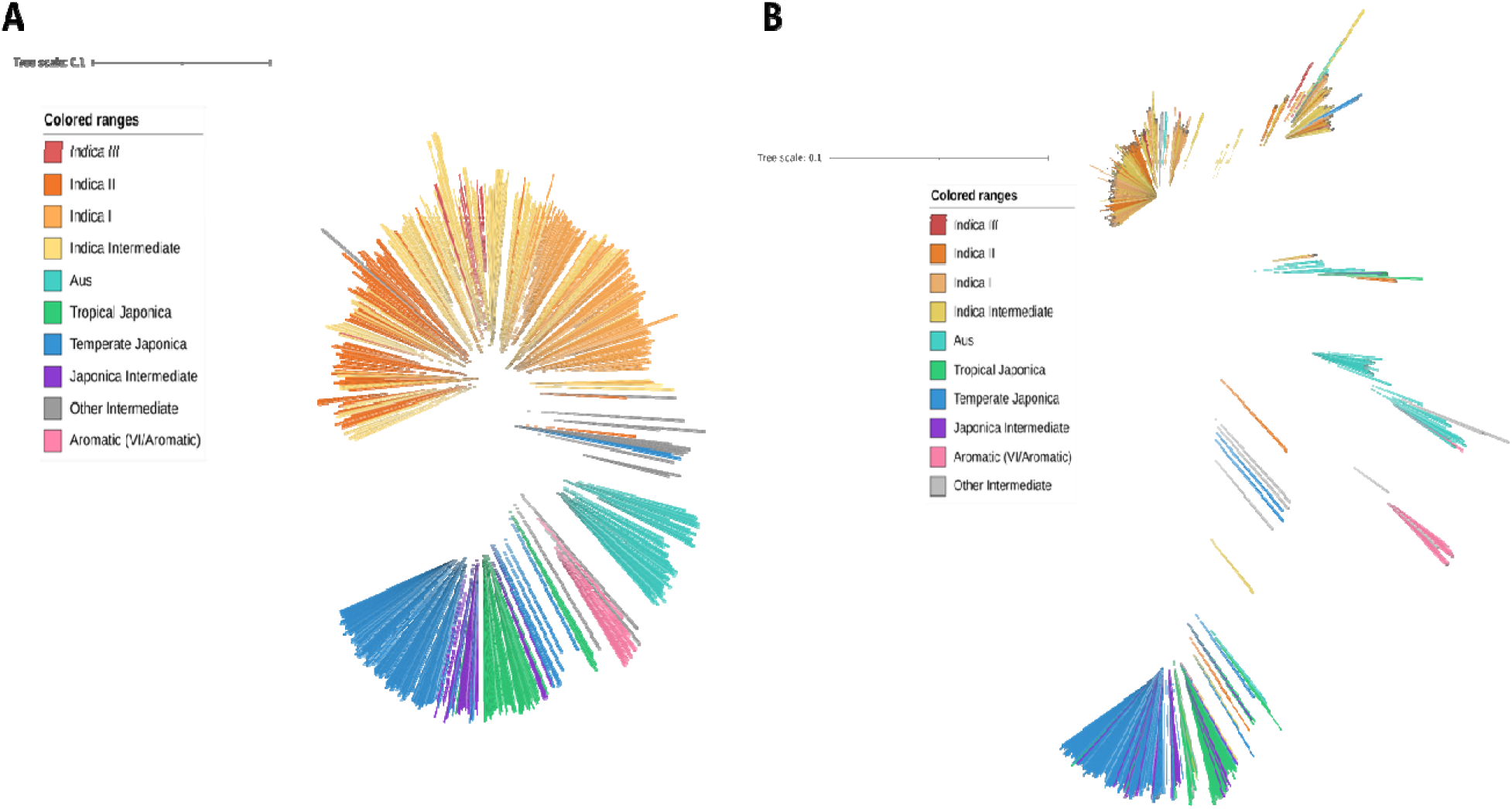
Genome-wide and IMI-specific phylogenetic relationships among rice accessions. Neighbor-joining trees constructed using SNPs from 533 cultivated rice accessions representing major Oryza sativa subpopulations. ***(A)*** shows the genome-wide phylogeny based on SNPs across the entire genome, whereas ***(B)*** shows the phylogeny constructed using SNPs restricted to the chromosome 6 IMI region. Branches are colored according to major rice subpopulation assignments as per the Rice 3k project, as indicated in the legend. While major subpopulation groupings are evident in both trees, differences in branch organization and clustering are apparent between the genome-wide and IMI-based phylogenies.

The IMI-based tree showed a similar overall grouping of major rice subpopulations. Within *indica*, accessions formed two distinct clusters in the IMI-based tree, whereas *indica* accessions were not separated into comparable subclusters in the genome-wide tree. The relative placement of non-*indica* groups was broadly consistent between the two trees.

Together, these results demonstrate that the chromosome 6 IMI region captures population structure that is not recapitulated by flanking regions or genome-wide variation of similar scale.

### Population genetic signatures across the chromosome 6 IMI region

To further characterize population genetic differentiation associated with the chromosome 6 IMI locus, nucleotide diversity (π), Tajima’s D, and genetic differentiation (FST) were estimated using four representative *indica* cultivars selected from the two *indica* clusters resolved specifically in the IMI-based PCA analysis (***Figure 8***). These cultivars were chosen because subdivision between the two groups was evident only when variants from the IMI region were analyzed and was absent in genome-wide or flanking-region PCA. Population genetic statistics were calculated in sliding windows across regions upstream of the IMI locus (pre-IMI), within the IMI interval, and downstream of the locus (post-IMI). In the pre-IMI region, nucleotide diversity values were comparable between the two *indica* groups, with π generally ranging between ∼0.0049 and 0.0050 across windows. Tajima’s D values fluctuated around neutral expectations, spanning approximately -1.0 to +1.5, while FST values remained low, typically below ∼0.2, indicating minimal genetic differentiation outside the IMI locus. In contrast, the IMI region exhibited a clear elevation in genetic differentiation. FST values increased substantially across the interval, rising from near-zero background levels in flanking regions to sustained values reaching ∼0.6-0.8 within the IMI locus. Nucleotide diversity remained broadly similar between the two groups (π ∼0.0049-0.0050), indicating that overall diversity was not reduced within the IMI interval. However, Tajima’s D profiles differed between the two groups across parts of the IMI region, with values spanning approximately -1.5 to +1.5 and showing pronounced divergence particularly toward the proximal portion of the locus, consistent with differences in allele frequency spectra rather than genome-wide demographic effects. In the post-IMI region, population genetic patterns again converged between the two *indica* groups. Nucleotide diversity remained within a similar range (π ∼0.0049-0.0050), Tajima’s D values overlapped broadly between groups (approximately -1.5 to +2.0), and FST values declined to lower levels, generally below ∼0.4, consistent with a return to a shared genomic background outside the structurally complex IMI interval.

**Figure 8.**
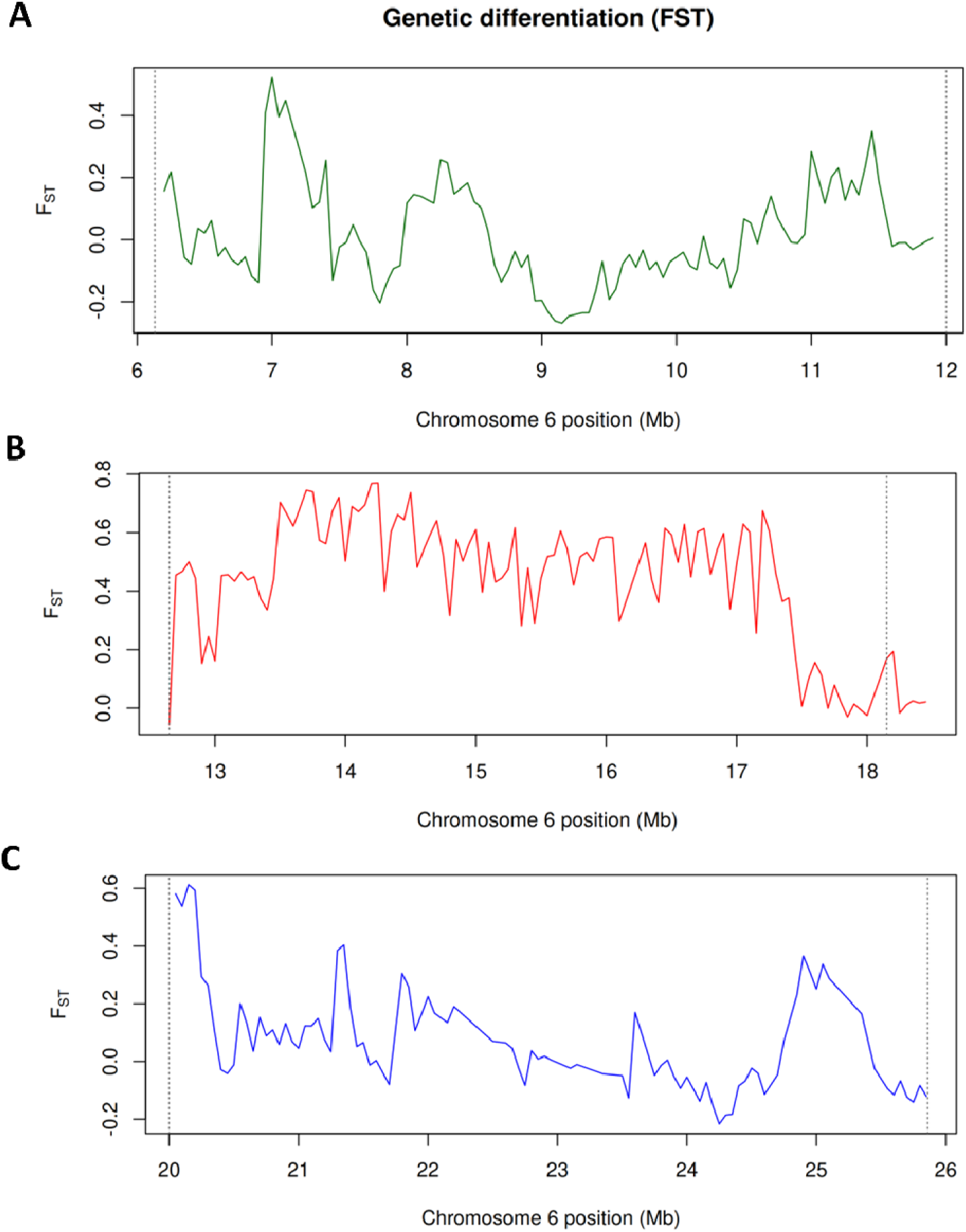
Genetic differentiation (FST) between two indica lineages across chromosome 6. Sliding-window estimates of genetic differentiation (FST) between two representative *indica* groups were calculated across chromosome 6 using variants aligned to the SM reference genome. Panels show ***(A)*** the region upstream of the IMI locus (pre-IMI), ***(B)*** the IMI interval, and ***(C)*** the downstream region (post-IMI). Vertical dashed lines indicate the boundaries of each analyzed interval. Elevated FST values are observed specifically within the IMI region, whereas flanking regions show comparatively lower differentiation, indicating that genetic divergence between the two *indica* groups is concentrated at the structurally complex IMI locus rather than distributed uniformly along the chromosome.

Together, these results demonstrate that genetic differentiation between the two *indica* clusters is quantitatively elevated in the IMI region, whereas flanking regions of chromosome 6 retain low differentiation and similar diversity levels between groups. No clear enrichment for geographic origin, breeding status, or *indica* subpopulation category was observed between the two IMI-defined clusters, indicating that the observed subdivision reflects structural haplotype segregation at this locus rather than demographic or geographic stratification.

## Discussion

### Genome architecture of Samba Mahsuri

The chromosome-scale assembly of *Oryza sativa* cv. SM provides a high-quality genomic reference from a predominantly *indica* genetic background. The assembled genome size of SM is comparable to other *indica* genomes, including R498 and 93-11, and larger than *japonica* reference genomes such as Nipponbare and Zhonghua 11, consistent with well-established subspecies-specific differences in repeat content (Du et al., 2017; Kawahara et al., 2013; Shand et al, 2023, Yao et al., 2025). Repetitive sequences account for ∼50.9% of the SM genome, a proportion closely matching that reported for *indica* references such as R498 and 93-11 and exceeding that of japonica genomes. As observed across rice genomes, repeats in SM are strongly enriched in pericentromeric and heterochromatic regions. Independent flow cytometric measurements further support this pattern, showing that *indica* rice possesses ∼9.7% higher nuclear DNA content than *japonica*, with genome size differences driven primarily by variation in repetitive sequence copy number rather than inflation of coding gene content (Ohmido et al., 2000).

Annotation of protein-coding genes in SM recovered a gene complement within the range reported for other high-quality rice genomes. Evidence-guided annotation predicted 31,138 protein-coding genes and 35,069 transcripts, with BUSCO completeness of 97.4%, comparable to or exceeding completeness reported for reference assemblies of Nipponbare, R498, and Zhonghua 11 (Du et al., 2017; Kawahara et al., 2013; Shand et al, 2023, Yao et al., 2025). As in other rice genome projects, apparent differences in gene counts across references likely reflect variation in assembly contiguity, repeat masking, transcript support, and annotation strategies rather than true lineage-specific gain or loss of core genes. The recovery of most conserved single-copy orthologs and the even chromosomal distribution of genes toward distal regions further support the completeness and biological plausibility of the SM gene annotation.

Non-coding RNA-coding gene annotation similarly indicates a high degree of completeness and conservation. The SM genome encodes 8,179 ncRNA loci, including 606 tRNA genes and the full complement of conserved rRNA, snoRNA, snRNA, and miRNA families expected in rice. As reported for Nipponbare and R498, most ncRNA families occur at low copy number and are broadly distributed, consistent with their conserved cellular roles. In contrast, repetitive ncRNA loci exhibit lineage-specific structural variation. Notably, 5S rDNA repeats form tandem arrays restricted to chromosome 11 in SM, Nipponbare, and R498, while differences in intergenic spacer length among these genomes reflect rapid, lineage-specific evolution of repetitive rDNA units under concerted evolution.

Annotation of organellar genomes further supports the canonical nature of the SM assembly. The SM chloroplast genome displays the typical quadripartite structure and gene content observed across *Oryza sativa*, with highly conserved gene order relative to Nipponbare and other rice cultivars (Du et al., 2017; Feng et al., 2021; Matsumoto, 2005). In contrast, the mitochondrial genome exhibits the structural plasticity characteristic of angiosperm mitochondria, with a conserved core gene set embedded within a dynamic architecture shaped by recombination, duplication, and fragmentation. In this context, differences in the presence or fragmentation of specific mitochondrial ORFs between SM and Nipponbare mirror patterns reported among other rice cultivars and align with evidence that plant mitochondrial genomes comprise heterogeneous, recombination-generated structures rather than single circles (Kozik et al., 2019).

Taken together, annotation-level comparisons indicate that the SM genome captures the conserved core features of rice nuclear and organellar genomes while exhibiting repeat-driven structural variation. Residual differences in predicted gene models or repeat organization are best interpreted in the context of assembly resolution and annotation sensitivity, particularly within highly repetitive regions. As a high-quality draft genome, SM therefore provides a reliable and complementary reference that expands the representation of genome diversity available for comparative and population genomic analyses.

### Genome-wide structural conservation and evolutionary context

Comparative analyses place the SM genome firmly within the canonical structural framework of rice genomes. At the chromosome scale, gene order and large-scale collinearity are strongly conserved relative to both the *japonica* reference Nipponbare and the *indica* reference R498, reflecting the stable macro-syntenic organization of rice chromosomes established following ancient whole-genome duplication events (Guyot & Keller, 2004; Wang et al., 2005). Much of the present-day rice genome comprises relics of these ancestral duplications, which have undergone extensive diploidization through gene loss while preserving overall chromosome structure (Wang et al., 2005). Consistent with this evolutionary history, duplicated genes retained in the SM genome predominantly show signatures of long-term purifying selection, a pattern widely observed among paleoduplicated genes in rice and other grasses and indicative of functional constraint rather than widespread neofunctionalization (Throude et al., 2009). Together, these observations indicate that the SM genome is structurally conservative at the whole-chromosome scale, shaped primarily by ancient duplication and subsequent diploidization rather than recent genome-wide innovation.

### Genome-wide divergence and heterogeneity

Genome-wide sequence and structural divergence between SM and the reference genomes is broadly distributed rather than concentrated in narrow genomic intervals. Divergence is more pronounced in comparisons between the predominantly *indica* SM genome and *japonica* reference genomes than in comparisons with canonical Chinese *indica* cultivar reference genomes (e.g. R498 and 93-11), reflecting lineage divergence rather than assembly artefacts or large-scale genome reorganization. Despite the presence of inversions, translocations, and non-aligned segments, large-scale synteny is largely preserved, and minor chromosome-specific length differences fall within the range reported for other rice assemblies, consistent with differential resolution of repeat-rich pericentromeric and heterochromatic regions (Du et al., 2017; Shang et al. 2023). Within this broadly conserved genomic framework, however, divergence is heterogeneous. While most of the genome retains strong macrosynteny, a subset of loci exhibits disproportionate structural complexity, duplication, or non-alignment. Similar patterns of focal structural heterogeneity have been reported among elite *indica* cultivars, including comparisons between Zhenshan 97 and Minghui 63, underscoring structural variation as a major axis of intrasubspecific diversity in rice (Zhang et al., 2016). The SM genome therefore contributes an additional, distinct reference, broadening the representation of genome diversity available for comparative analyses.

### The chromosome 6 IMI locus represents an extreme example of locus-specific structural complexity

One of the most striking focal regions in our comparative genomic study using the SM genome is the chromosome 6 IMI locus. This region corresponds to a large pericentromeric interval previously reported as a single inversion in the R498 genome and later described as a nested inversion structure in 93-11 (Du et al. 2017; Zhang et al. 2016). However, prior studies treated this region primarily as an isolated structural feature and did not investigate its population-level consequences.

Using SM as the reference, we show that the IMI locus has a direct match in the *indica* reference R498, contrasting sharply with the configuration observed in the *japonica* reference Nipponbare. Comparative analyses of wild *Oryza* genomes further indicate that multiple orientations and nested rearrangements segregate at this locus, pointing to an evolutionary dynamic region, of pre-domestication origin. Extending this analysis through joint genotyping of 533 cultivated rice accessions reveals that IMI configurations are strongly associated with major rice lineages, yet remain structurally heterogeneous, with multiple orientations and nested rearrangements segregating within and across subspecies. These patterns indicate that IMI represents a lineage-biased but evolutionarily dynamic locus, rather than a single fixed rearrangement confined to one genetic background. Why does the IMI locus generate population structure that is invisible elsewhere? Population-scale analyses of 533 rice cultivars reveal that genetic variation within the IMI region produces a distinct population structure that is absent in flanking regions and at the whole-chromosome level. PCA based on IMI variants resolves two clusters within *indica* rice, a pattern that collapses when variants from adjacent regions, whole chromosome 6, or chromosome 1 are analyzed. Importantly, well-studied *indica* genomes such as Zhenshan 97 and Minghui 63 fall into different IMI-defined clusters, consistent with their known structural divergence despite shared subspecies identity (Zhang et al., 2016). Notably, nucleotide diversity and Tajima’s D do not show pronounced IMI-specific deviations relative to flanking regions, arguing against recent selective sweeps or genome-wide demographic effects as drivers of differentiation. Instead, the strong elevation of FST confined to the IMI interval supports a model in which alternative structural haplotypes are maintained by reduced recombination, generating population structure without substantially altering local allele-frequency spectra.

### Implications: structural polymorphism as a driver of locus-specific evolutionary dynamics

Large inversions and nested rearrangements suppress recombination, allowing divergent haplotypes to persist over long timescales despite ongoing gene flow elsewhere in the genome (Crow et al., 2020). Consistent with this expectation, our analysis indicates that the chromosome 6 IMI region follows an evolutionary trajectory that is partially decoupled from the rest of the genome. Although the IMI structure has been described previously, its population-level consequences have not been resolved.

The IMI-defined haplotype clusters within rice do not show clear geographic or ecological segregation, suggesting that the observed structure is not primarily driven by regional adaptation. Moreover, while inversions can contribute to adaptive divergence in some systems, we do not directly assess phenotypic or functional effects here. The observed clustering is therefore best interpreted as reflecting long-standing structural polymorphism rather than evidence of recent positive selection. Experimental studies in rice have shown that large inversions strongly suppress recombination in indica crosses (Zhou et al., 2023), providing a plausible mechanistic basis for the persistence of divergent IMI haplotypes and the locus-specific population structure detected in this study.

More broadly, large inversion polymorphisms remain challenging to study because reduced recombination limits fine-mapping, ancestry correlations complicate association analyses, and the large number of genes involved obscures the identification of causal variants for different traits Nonetheless, our results demonstrate that the IMI region functions as a distinct evolutionary unit within the rice genome. By anchoring population-scale analyses to an appropriate reference genome, we reveal strong, locus-specific population structure that is invisible in genome-wide analyses. These findings underscore how large structural variants can locally shape genetic diversity and highlight the importance of genome structure in population genomic inference.

## Data availability

Raw sequencing reads and the CSIR-CCMB-SMv1.0 genome assembly have been deposited at NCBI under BioProject PRJNA1242182.

## Author Contributions

DR, RVS, ST, and HKP conceived and designed the experiments. DR and SKK contributed equally to this work. DR performed plant growth, tissue collection for DNA and RNA sequencing, and nucleic acid isolation; carried out chromosome-scale genome assembly, ab initio gene prediction, and functional annotation; conducted comparative genomics and evolutionary analyses; discovered and characterized the chromosome 6 IMI locus; and wrote the first draft of the manuscript. SKK performed evidence-guided gene prediction and functional annotation, organelle genome assembly and annotation, ncRNA annotation, developed the SnpEff database for the SM genome, and conducted population genomic analyses. NT maintained plant material, collected tissues for RNA sequencing at different developmental stages, and assisted with RNA isolation. EK contributed to variant analyses. HKP, ST, and RVS supervised the research, provided conceptual guidance, edited the manuscript and acquired funding. All authors read and approved the final manuscript.

## Supporting information

Supplementary sheet

## Acknowledgements

We thank Dr. Sheshu Madhav Maganti (formerly at IIRR; currently at NIRCA) for facilitating the procurement of seed material from the nucleus seed stock of *Oryza sativa* cv. Samba Mahsuri at Agricultural Research Station (ARS), Bapatla. We also thank Dr. Raghunand R. Tirumalai (CSIR-CCMB) for helpful discussions as a member of the doctoral advisory committee of DR, including inputs on the IMI locus. We thank Dr. C G Gokulan for thought-provoking discussions. We acknowledge the sequencing facility at CSIR-CCMB for Illumina sequencing, and Nucleome Informatics Pvt. Ltd. for PacBio sequencing and Bionano optical mapping data generation. Additionally, we thank Dr. Raju Madanala and Dr. Jamaloddin for their assistance with indenting and maintenance of the rice fields, respectively. We also thank our IT team for supporting us with the maintenance of our servers for data analysis. We acknowledge the RiceVarMap2 and 3,000 Rice Genomes Project consortia for generating and making publicly available population-scale rice sequencing data used in this study.

## Abbreviations and full forms

SM-: Samba Mahsuri (*Oryza sativa* cv. Samba Mahsuri)
NCBI-: National Center for Biotechnology Information
SRA-: Sequence Read Archive
MAF-: Minor Allele Frequency
SNP-: Single-Nucleotide Polymorphism
InDels-: Insertions Deletions
PCA-: Principal Component Analysis
IBS-: Identity-by-State
LD-: Linkage Disequilibrium
IMI-: Inversion-Match-Inversion
BUSCO-: Benchmarking Universal Single-Copy Orthologs
DLE-1-: Direct Label and Enzyme 1 (Bionano optical mapping chemistry)
LSC-: Large Single-Copy region (plastid genome terminology)
SSC-: Small Single-Copy region (plastid genome terminology)
IR-: Inverted Repeat (plastid genome terminology)
CCS-: Circular Consensus Sequencing (PacBio HiFi reads)
LTR-: Long Terminal Repeat
GO-: Gene Ontology
KEGG-: Kyoto Encyclopedia of Genes and Genomes
ORF-: Open Reading Frame
MYA-: Million Years Ago

