## Supplementary figures for "Genome Assembly of the Iconic Samba Mahsuri Delineates Locus-specific Population Structure within *Indica* Rice"

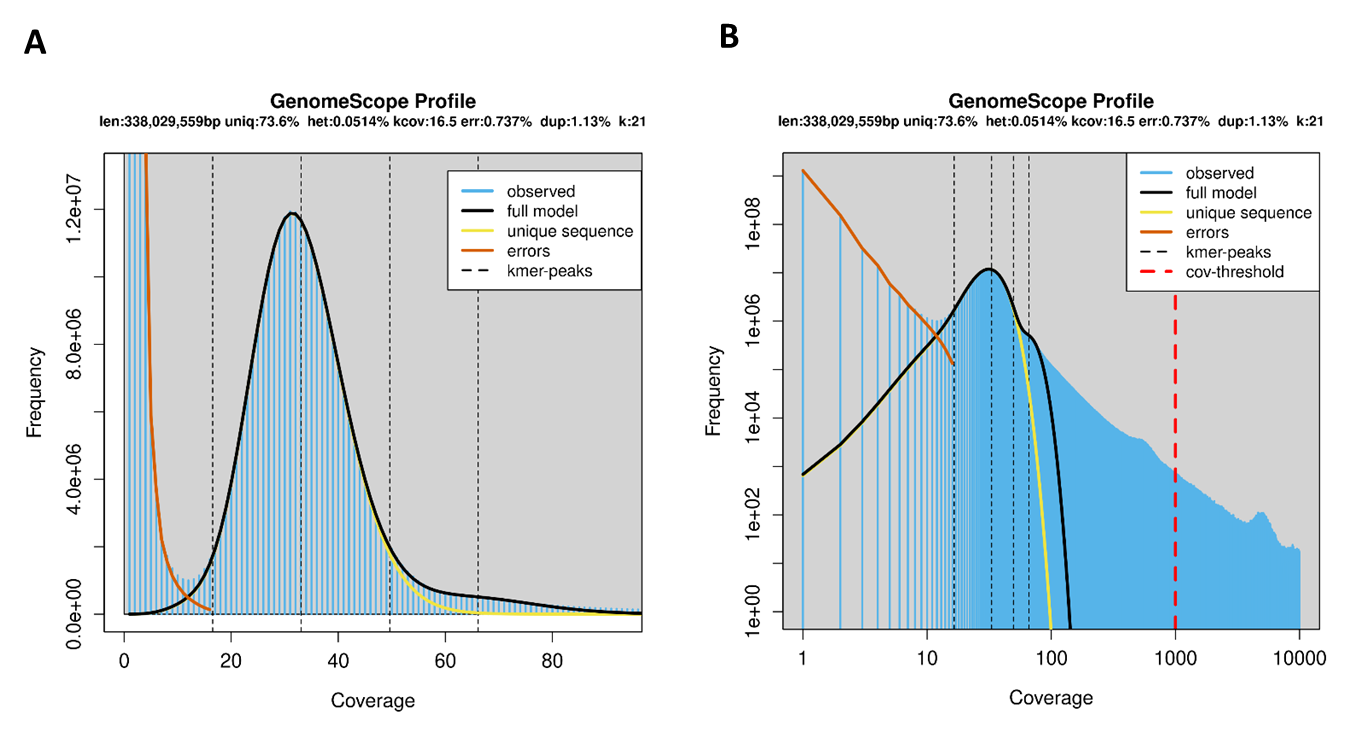

Figure S1: GenomeScope k-mer profile of the Samba Mahsuri (SM) genome. (A) K-mer frequency distribution (k = 21). The estimated haploid genome length is ~338 Mb, with ~0.05% heterozygosity, ~0.74% read-error rate, and ~73.6% unique sequence content. (B) Log-scale representation of the same profile illustrating the coverage distribution across unique and repetitive k-mers. The red dashed line marks the high-coverage cutoff used to exclude extreme repetitive k-mers.

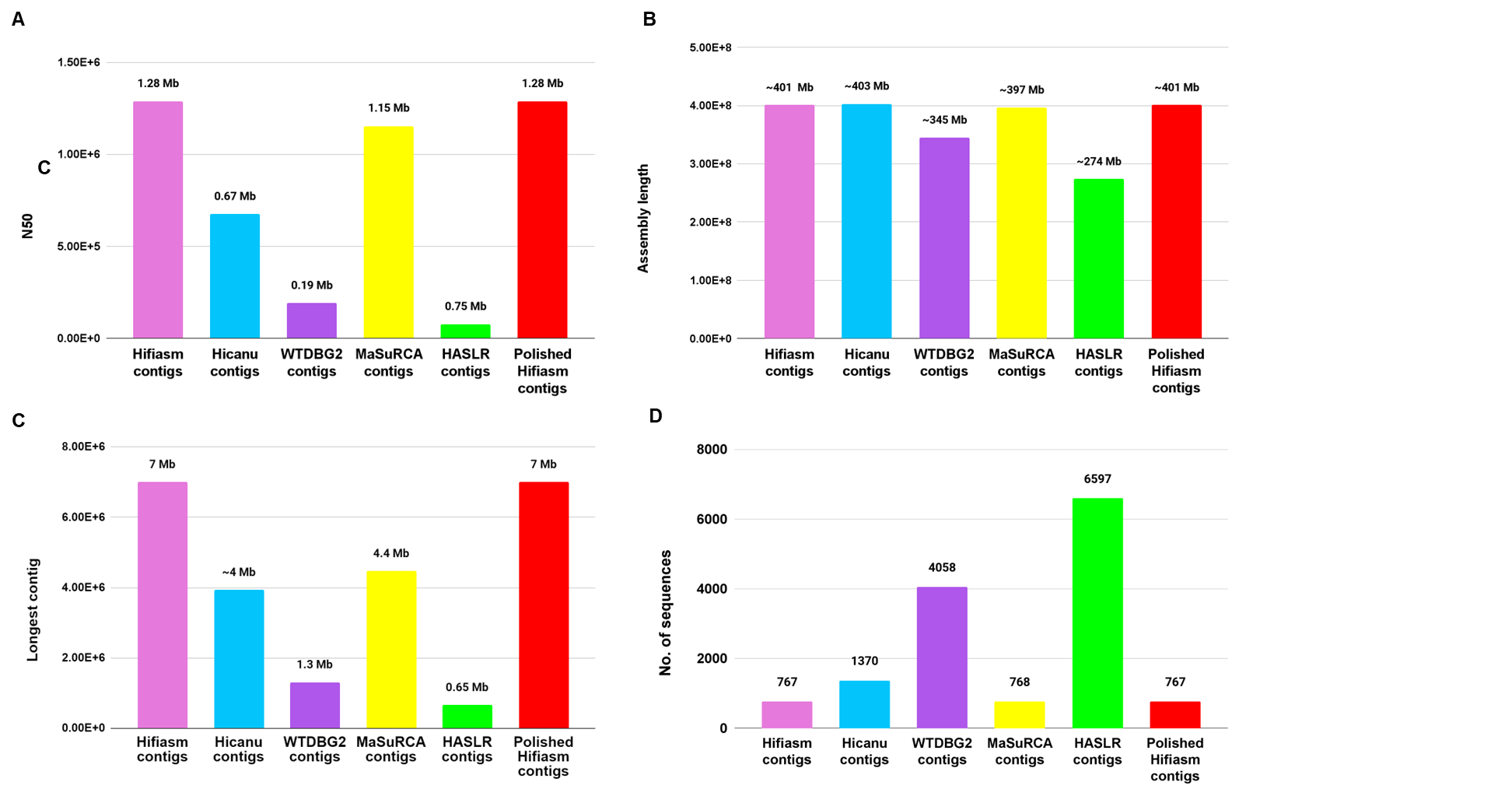

**Figure S2: Comparison of assembly quality metrics across different contig sets generated using multiple assembly strategies.** *Bar plots show* ***(A)*** *N50 values,* ***(B)*** *total assembly length,* ***(C)*** *length of the longest contig, and* ***(D)*** *number of sequences (contigs) in Hifiasm, Hicanu, WTDBG2, MaSuRCA, HASLR, and polished Hifiasm assemblies. Overall, the Hifiasm assembly exhibits higher contiguity (larger N50 and longest contig lengths) and lower fragmentation compared to other assemblers, while HASLR and WTDBG2 show more fragmented assemblies with a larger number of contigs. Hence, it was polished and used as the final contig assembly.*

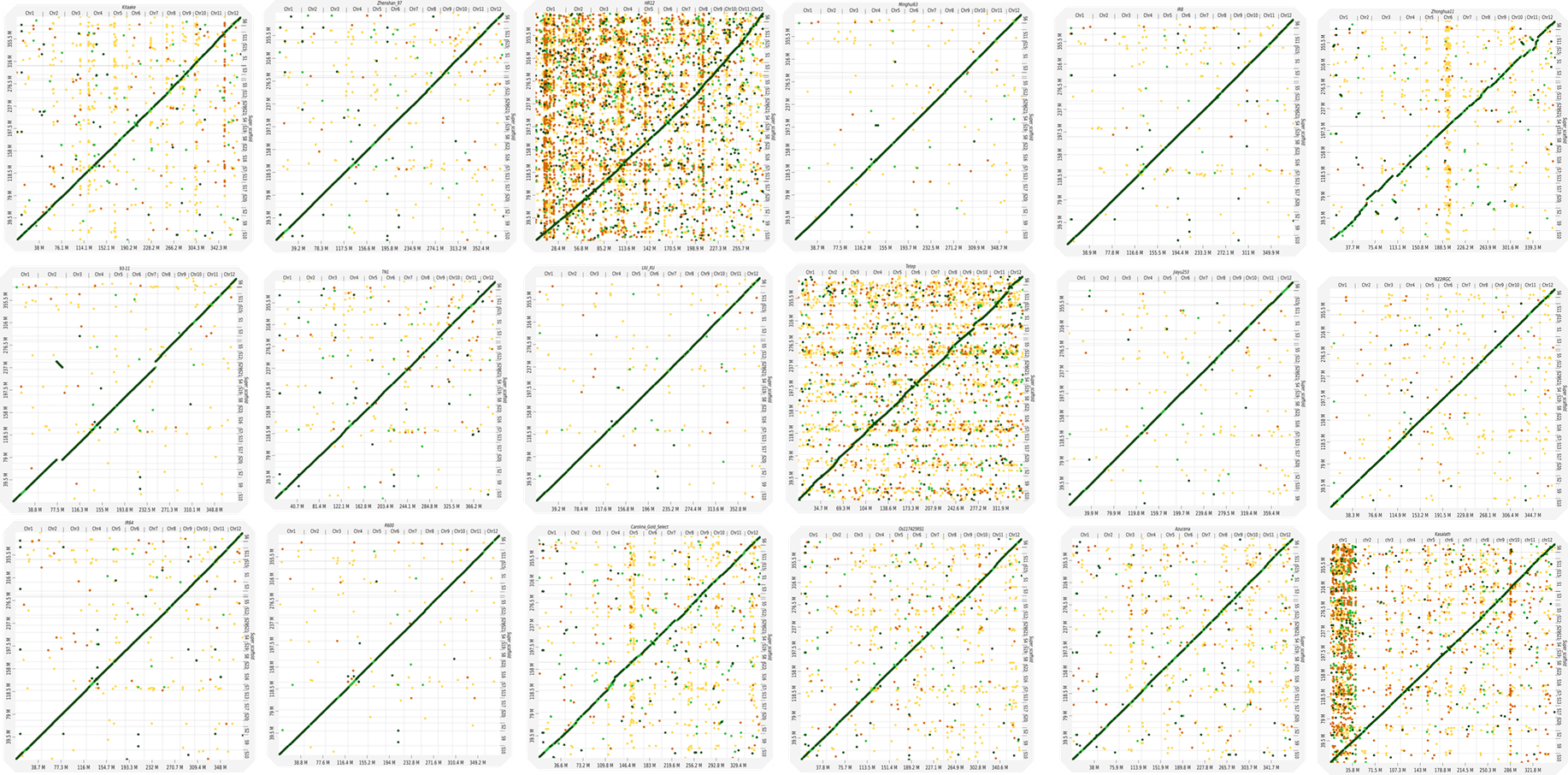

Figure S3: Chromosome-level dot plot comparisons between the Samba Mahsuri (SM) genome and all Oryza sativa assemblies available at chromosome scale in NCBI. They reveal strong genome-wide collinearity, with continuous diagonals indicating conserved synteny and scattered off-diagonal signals reflecting local structural variation. These comparisons were used to validate scaffold ordering and orientation of the SM assembly after polishing and BioNano scaffolding, with final orientations consistent with all Oryza sativa genomes.

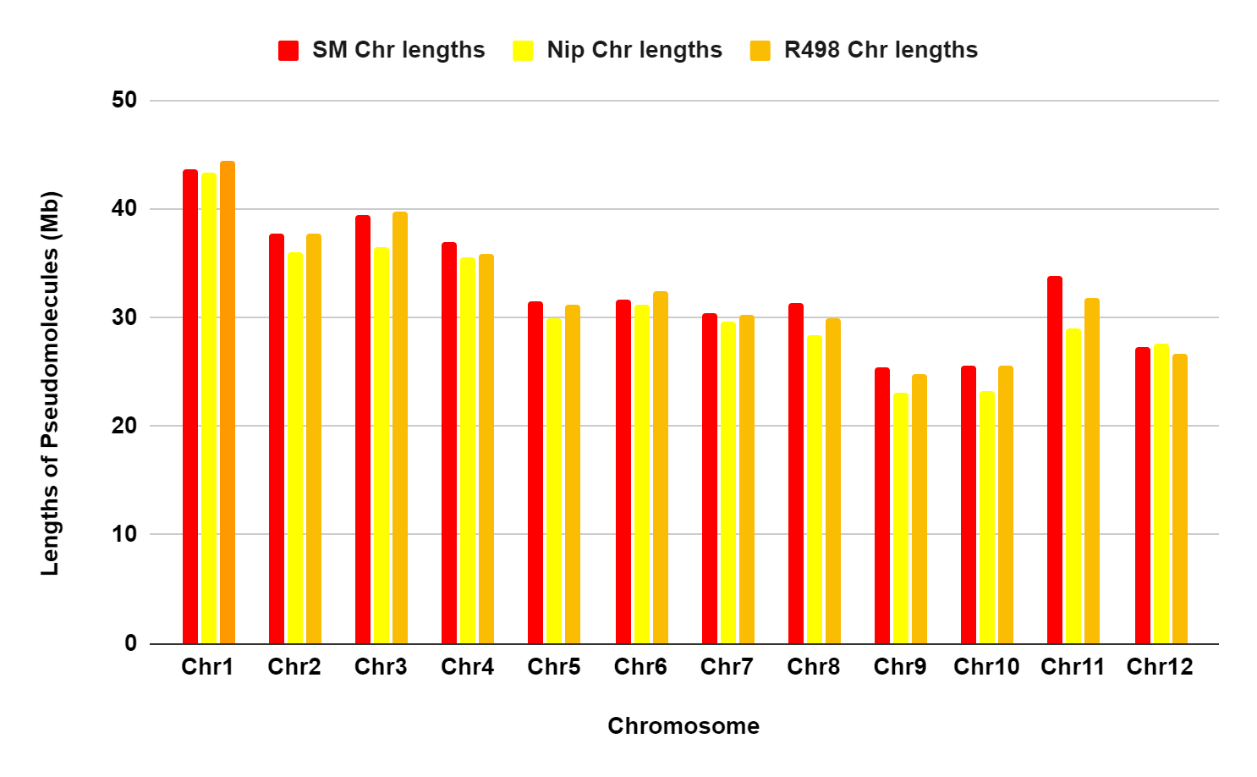

***Figure S4:*** ***Bar plots show the lengths of the 12 pseudomolecules in the Samba Mahsuri (SM) assembly compared with the corresponding chromosomes of the Nipponbare (IRGSP-1.0) and R498 genomes.*** SM chromosomes generally fall within the size range observed in the two reference genomes, with expansions notable in chromosomes 4, 5, 7, 8, 9, and 11. R498 exhibits the longest chromosomes for Chr1, Chr2, Chr3, Chr6, and Chr10, whereas Nipponbare displays slightly longer Chr12. Total pseudomolecule lengths are 394.7 Mb for SM, 390.3 Mb for R498, and 373.2 Mb for Nipponbare, indicating that SM is the largest among the three assemblies.

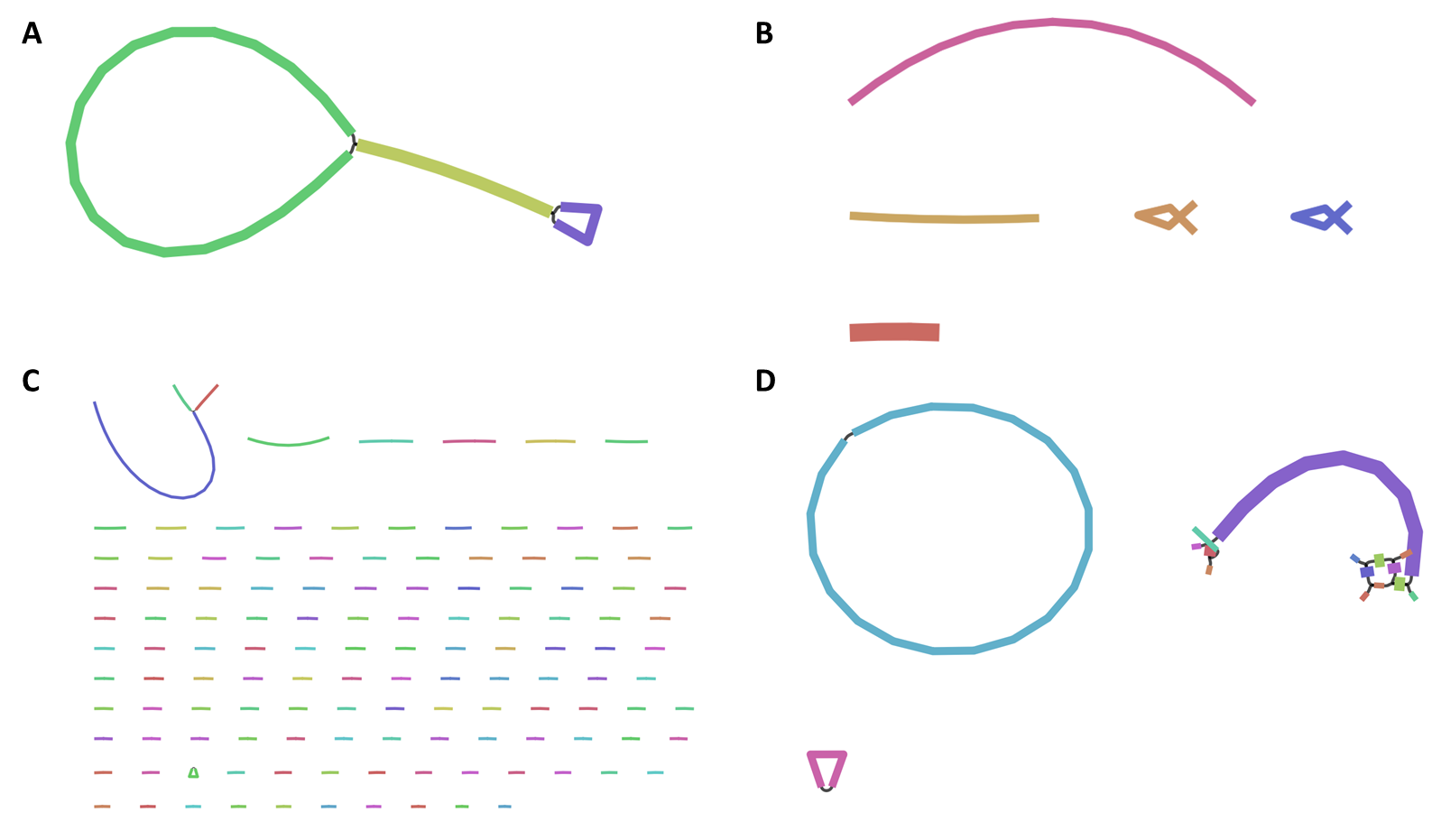

Figure S5: Organelle genome assemblies of Samba Mahsuri (SM) using multiple reconstruction methods. (A) The complete plastome assembly generated using GetOrganelle, showing a canonical circular structure with a well-resolved large single-copy (LSC), small single-copy (SSC), and inverted repeat (IR) configuration. (B) Mitochondrial genome assembly produced with GetOrganelle, consisting of several contigs and sub-genomic circles typical of plant mitochondrial complexity. (C) Mitochondrial genome reconstructed using MitoHifi, resulting in a single dominant circular mitochondrial chromosome with additional low-abundance alternative isoforms. (D) Assembly of the mitochondrial genome using Unicycler, revealing multiple interconnected circular and branched structures, consistent with known multipartite plant mitogenome architectures.

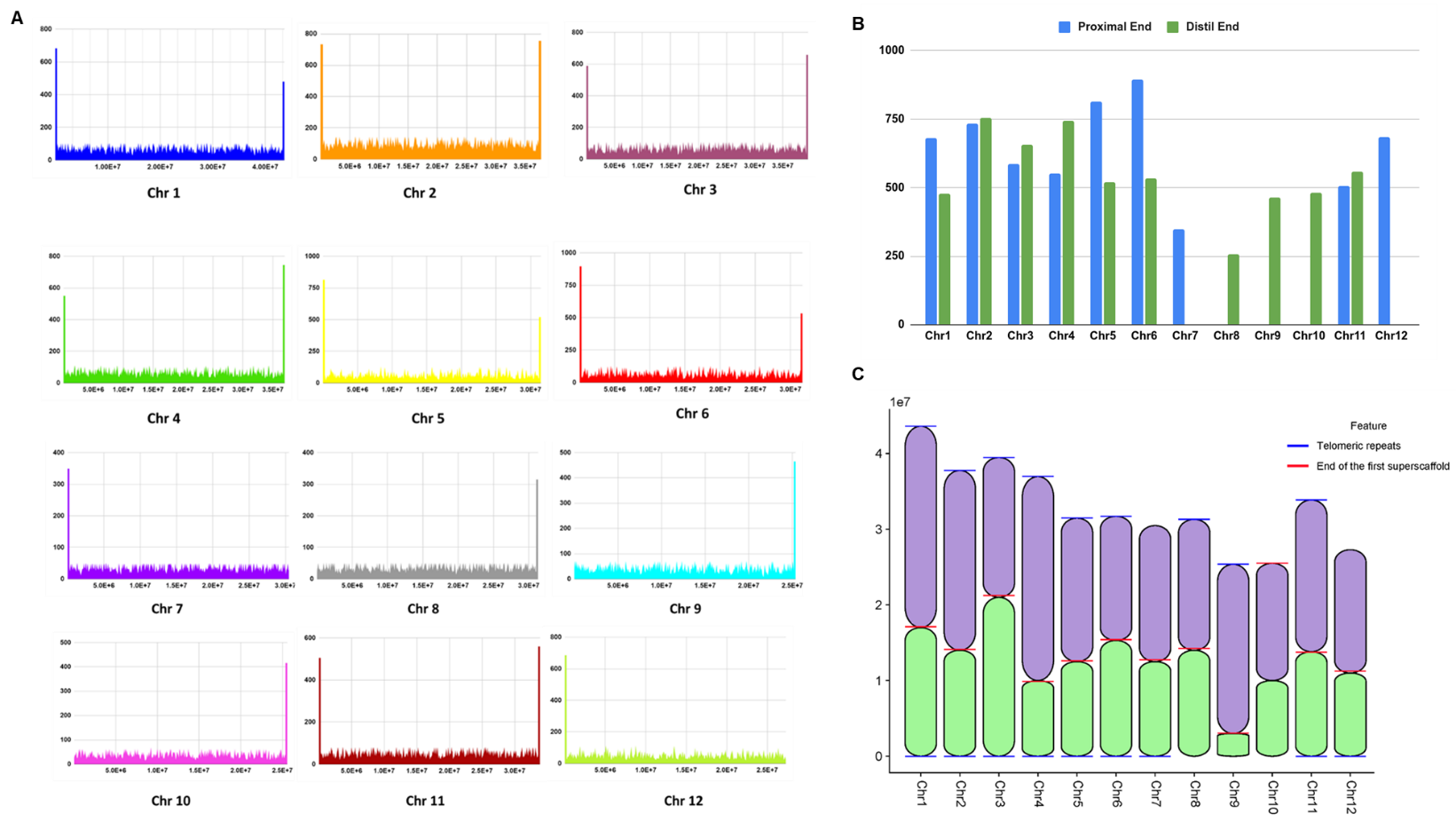

***Figure S6: Telomeres and centromeres in Samba Mahsuri (SM) pseudomolecules. (A)*** Telomeric repeat arrays (AAACCCT)n were detected at both ends of Chr1-6 and Chr11; at the proximal ends of Chr7 and Chr12; and at the distal ends of Chr8-10. ***(B)*** The bar chart shows the number of telomeric repeats at the ends of SM pseudomolecules. ***(C)*** Blue bars mark the positions of telomeric repeat arrays (AAACCCT)n. Red bars indicate the end of the first superscaffold used to assemble each chromosome. Most chromosomes were assembled from 2-3 superscaffolds. In most chromosomes, the superscaffold break coincides with the centromeric region, indicating that the assembly was interrupted at the highly repetitive RCS2-rich centromeres. These breaks were subsequently resolved and joined by anchoring against reference genomes. Notably, Chr10 was assembled as a single superscaffold without any internal break, indicating complete continuity across both arms.

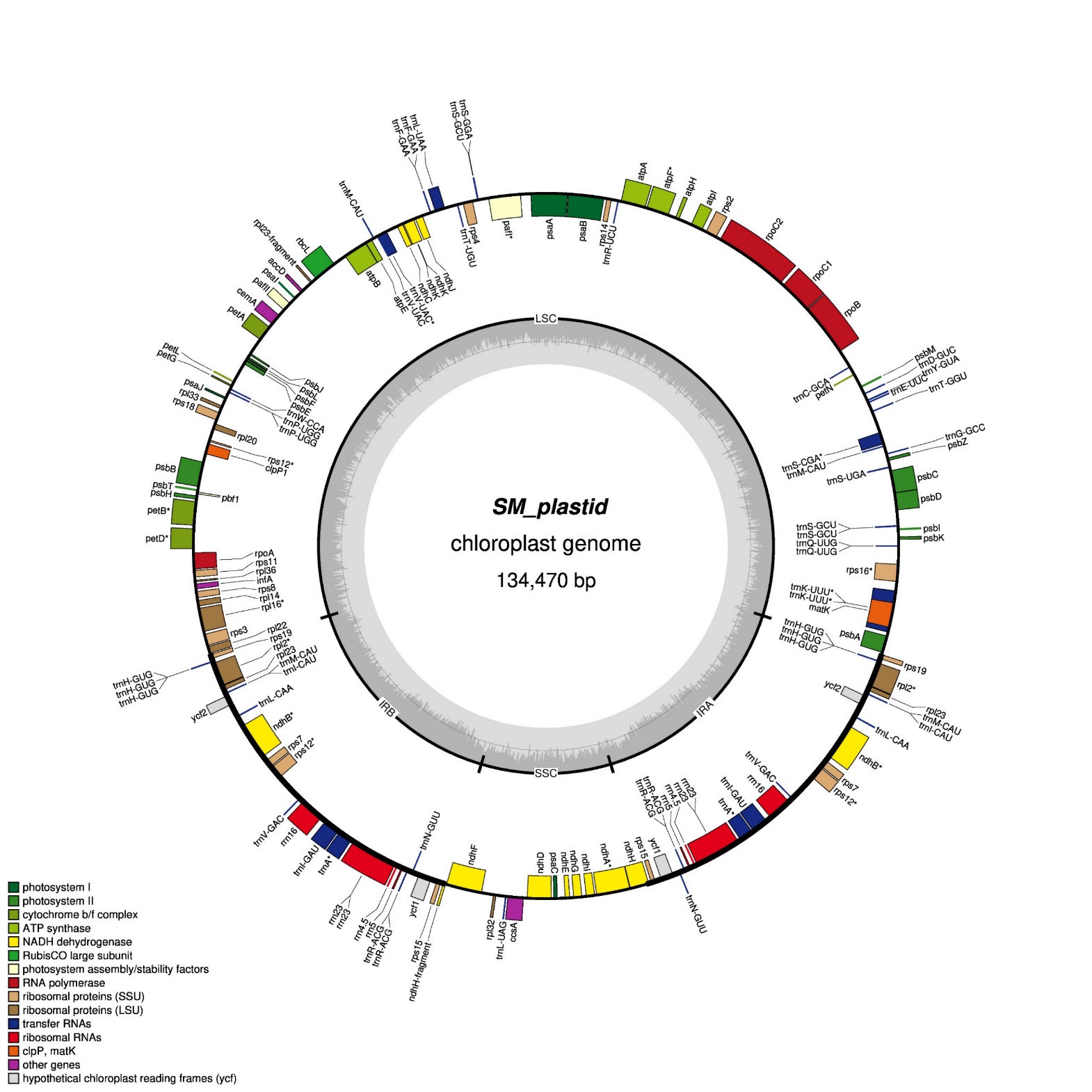

Figure S7: Annotated chloroplast genome of Samba Mahsuri (SM). The complete plastid genome of SM is shown as a circular map of 134,470 bp, displaying the canonical quadripartite structure comprising a large single-copy (LSC) region, a small single-copy (SSC) region, and two inverted repeats (IRa and IRb). Genes are color-coded by functional category, including photosystem I and II components, cytochrome b/f complex subunits, ATP synthase, NADH dehydrogenase, Rubisco large subunit, RNA polymerase, ribosomal proteins (LSU and SSU), transfer RNAs, ribosomal RNAs, and hypothetical open reading frames (ycf). Genes located inside the circle are transcribed clockwise, and those outside are transcribed counterclockwise. The inner rings show GC content and GC skew across the plastome.

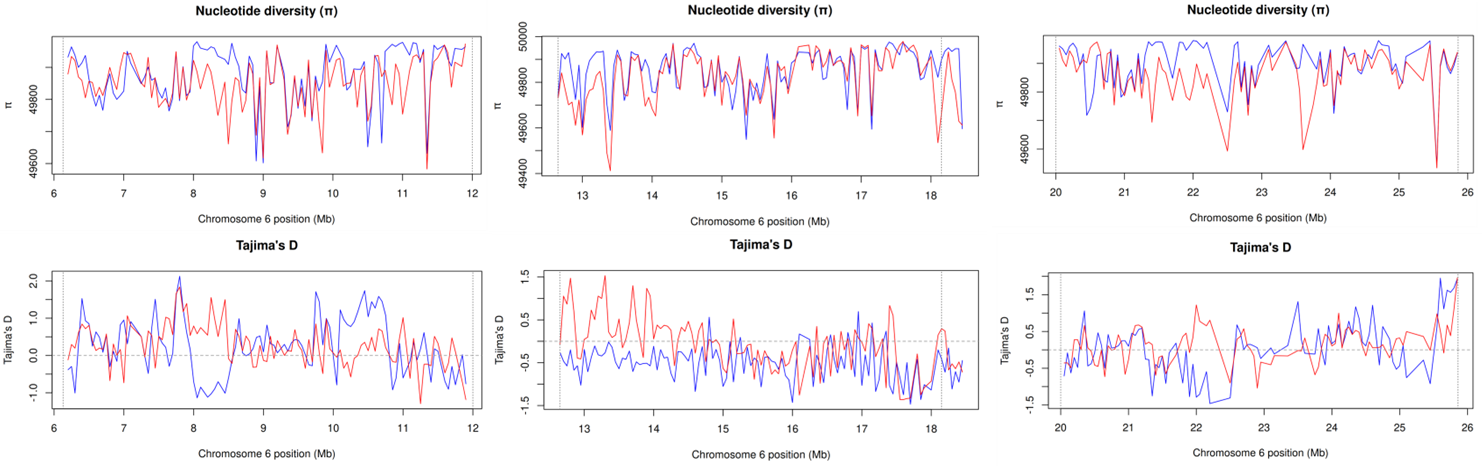

Figure S8: Population genetic signatures across the chromosome 6 IMI locus. Sliding-window profiles of nucleotide diversity (π), and Tajima’s D across pre-IMI, IMI, and post-IMI regions of chromosome 6 reveal distinct diversity and differentiation patterns within the inversion relative to flanking intervals.
