## Supplementary table for "Genome Assembly of the Iconic Samba Mahsuri Delineates Locus-specific Population Structure within *Indica* Rice"

Table S1: Comparison of hybrid and long-read genome assemblies of Samba Mahsuri (SM). Summary of key contiguity and completeness metrics obtained using different assemblers with PacBio CCS reads alone or in combination with Illumina short reads. Hifiasm produced the most contiguous and complete assembly, which was subsequently polished with Pilon using Illumina data to obtain the final reference assembly.

| **Assembler** | **Type** | **Contigs** | **Total Assembly Length (bp)** | **Mean Length (bp)** | **Longest Contig (bp)** | **Shortest Contig (bp)** | **N50 (bp)** | **L50** | **BUSCO (C[S,D], F, M)** |
| --- | --- | --- | --- | --- | --- | --- | --- | --- | --- |
| HASLR | Hybrid (PacBio + Illumina) | 6,602 | 273,802,680 | 41,473 | 654,579 | 483 | 75,194 | 1,036 | C:94.9% [S:94.3%, D:0.6%], F:0.6%, M:4.5% |
| MaSuRCA | Hybrid (PacBio + Illumina) | 768 | 396,712,634 | 516,553 | 4,467,369 | 5,396 | 1,152,959 | 103 | C:97.8% [S:97.2%, D:0.6%], F:0.5%, M:1.7% |
| HiCanu | PacBio CCS | 1,370 | 402,993,104 | 294,156 | 3,932,143 | 10,780 | 673,753 | 180 | C:97.0% [S:95.8%, D:1.2%], F:0.6%, M:2.4% |
| Wtdbg2 | PacBio CCS | 4,058 | 346,851,338 | 85,473 | 1,315,833 | 4,080 | 191,399 | 506 | C:96.6% [S:95.8%, D:0.8%], F:1.0%, M:2.4% |
| Hifiasm | PacBio CCS | 767 | 400,996,022 | 522,811 | 7,003,306 | 10,142 | 1,286,932 | 94 | C:97.3% [S:96.3%, D:1.0%], F:0.7%, M:2.0% |
| Hifiasm polished with Pilon | PacBio + Illumina | 767 | 400,940,681 | 522,739 | 7,001,649 | 10,141 | 1,286,484 | 94 | C:97.9% [S:96.9%, D:1.0%], F:0.4%, M:1.7% |
| Pilon Hifiasm + Optical mapping | PacBio + Illumina + Bionano | 30 | 394,983,264 | 13,166,109 | 27,061,646 | 73,873 | 16,300,341 | 10 | C:97.7%  [S:96.9%,  D:0.8%],  F:0.3%,  M:2.0% |
| Ordered pseudomolecule assembly | Ordered pseudomolecules (reference-guided) | 12 | 394,985,064 | 32,915,422 | 43,791,869 | 25,369,104 | 31,699,651 | 6 | C:97.6% [S:96.8%, D:0.8%], F:0.3%, M:2.1% |

Table S2: Chromosome Lengths of Samba Mahsuri (SM)

| **S. No.** | **Chromosome** | **Super-scaffolds** | **Length (bp)** | **SM Lengths (bp)** |
| --- | --- | --- | --- | --- |
| 1. | Chr1 | S10 | 17121206 | 43614007 |
|  |  | S9 | 26492801 |  |
| 2. | Chr2 | S2 | 14124282 | 37756426 |
|  |  | S14 | 10678929 |  |
|  |  | S20 | 12953215 |  |
| 3. | Chr3 | S17 | 21235636 | 39466463 |
|  |  | S13 | 18230827 |  |
| 4. | Chr4 | S7 | 9897847 | 36959493 |
|  |  | S16 | 27061646 |  |
| 5. | Chr5 | S22 | 12617328 | 31463684 |
|  |  | S8 | 18846356 |  |
| 6. | Chr6 | S19 | 15399210 | 31699551 |
|  |  | S4 | 16300341 |  |
| 7. | Chr7 | S21 | 12759277 | 30470816 |
|  |  | S29 | 12653262 |  |
|  |  | S18 | 5058277 |  |
| 8. | Chr8 | S12 | 14237405 | 31292396 |
|  |  | S5 | 17054991 |  |
| 9. | Chr9 | S24 | 3058987 | 25368904 |
|  |  | S30 | 6916040 |  |
|  |  | S3 | 15393877 |  |
| 10. | Chr10 | S1 | 25496796 | 25496796 |
| 11. | Chr11 | S15 | 13741604 | 33856046 |
|  |  | S25 | 354384 |  |
|  |  | S11 | 19760058 |  |
| 12. | Chr12 | S23 | 11245481 | 27287147 |
|  |  | S27 | 197450 |  |
|  |  | S6 | 15844216 |  |

Table S3: Comparative lengths of chromosomes/pseudomolecules in Samba Mahsuri (SM), Nipponbare, and R498 rice varieties. The table lists chromosome-wise lengths in base pairs (bp) for SM, Nipponbare, and R498, along with a summary of relative length comparisons across the three varieties.

| **S. No.** | **Chromosome** | **SM Chr lengths**  **(bp)** | **Nipponbare Chr lengths**  **(bp)** | **R498 Chr lengths**  **(bp)** | **Length comparisons** |
| --- | --- | --- | --- | --- | --- |
| 1. | Chr1 | 43,614,007 | 43,270,923 | 44,361,539 | R498>SM>Nip |
| 2. | Chr2 | 37,756,426 | 35,937,250 | 37,764,328 | R498>SM>Nip |
| 3. | Chr3 | 39,466,463 | 36,413,819 | 39,691,490 | R498>SM>Nip |
| 4. | Chr4 | 36,959,493 | 35,502,694 | 35,849,732 | SM>R498>Nip |
| 5. | Chr5 | 31,463,684 | 29,958,434 | 31,237,231 | SM>R498>Nip |
| 6. | Chr6 | 31,699,551 | 31,248,787 | 32,465,040 | R498>SM>Nip |
| 7. | Chr7 | 30,470,816 | 29,697,621 | 30,277,827 | SM>R498>Nip |
| 8. | Chr8 | 31,292,396 | 28,443,022 | 29,952,003 | SM>R498>Nip |
| 9. | Chr9 | 25,368,904 | 23,012,720 | 24,760,661 | SM>R498>Nip |
| 10. | Chr10 | 25,496,796 | 23,207,287 | 25,582,588 | R498>SM>Nip |
| 11. | Chr11 | 33,856,046 | 29,021,106 | 31,778,392 | SM>R498>Nip |
| 12. | Chr12 | 27,287,147 | 27,531,856 | 26,601,357 | Nip>SM>R498 |
| 13. | **Total** | **394,731,729** | **373,245,519** | **390,322,188** | **SM>R498>Nip** |

Table S4: RCS2 Family of Repeats in the Samba Mahsuri (SM) Genome

| **S. No.** | **Chromosome** | **RCS2 Repeat Location** | **Probable Centromeric Location** |
| --- | --- | --- | --- |
| 1. | Chr1 | 17069755-17302181 | ~17 Mb |
| 2. | Chr2 | 14079007-14140477 | ~14 Mb |
| 3. | Chr3 | 21152640-21302859,  29304-29402 | ~21 Mb |
| 4. | Chr4 | 9876428-9908192 | ~9-10 Mb |
| 5. | Chr5 | 12599079-12640384 | ~12.5 Mb |
| 6. | Chr6 | 15392720-15401425 | ~15.3 Mb |
| 7. | Chr7 | 12577214-12774787 | ~12.5 Mb |
| 8. | Chr8 | 14185094-15147994,  29606580-29606736 | ~14-15 Mb |
| 9. | Chr9 | 2615836-2652299,  3051678-3066335,  3114512-3115494,  3227100-3229058 | ~2.6-3.2 Mb |
| 10. | Chr10 | 9499363-10595388 | ~9-10.5 Mb |
| 11. | Chr11 | 13473466-14105177 | ~13-14 Mb |
| 12. | Chr12 | 11202451-11443944 | ~11 Mb |

Table S5: Classification and genome coverage of repetitive elements in the Samba Mahsuri (SM) genome

| Repeat Type |  |  |  | Repeat Element | Length | % of Genome Length |
| --- | --- | --- | --- | --- | --- | --- |
| Retroelements: |  |  |  | 80577 | 95776920 | 24.26 |
|  | LINEs: |  |  | 9064 | 4195567 | 1.06 |
|  |  | RTE/Bov-B |  | 154 | 99855 | 0.03 |
|  |  | L1/CIN4 |  | 8910 | 4095712 | 1.04 |
|  | LTR elements: |  |  | 71513 | 91581353 | 23.2 |
|  |  | BEL/Pao |  | 277 | 60250 | 0.02 |
|  |  | Ty1/Copia |  | 13727 | 11376471 | 2.88 |
|  |  | Gypsy/DIRS1: |  | 29686 | 67933537 | 17.21 |
|  |  |  | Retroviral | 3616 | 1055818 | 0.27 |
| DNA transposons: |  |  |  | 18332 | 14617748 | 3.7 |
|  | hobo-Activator |  |  | 3972 | 2304459 | 0.58 |
|  | Tourist/Harbinger |  |  | 805 | 469438 | 0.12 |
| Rolling-circles |  |  |  | 1653 | 1075571 | 0.27 |
| Unclassified |  |  |  | 354736 | 83995719 | 21.28 |
| Total interspersed repeats |  |  |  |  | 195465958 | 49.24 |
| Small RNA |  |  |  | 1134 | 434124 | 0.11 |
| Satellites |  |  |  | 411 | 209585 | 0.05 |
| Simple repeats |  |  |  | 93203 | 4366108 | 1.11 |
| Low complexity |  |  |  | 9993 | 500675 | 0.13 |
| Total |  |  |  | 560039 | 200976450 | 50.9 |

Table S6: Gene Prediction Dataset Metric Comparison

| **Dataset Metrics** | **Evidence-based Transcripts** | **Genemark Predictions** | **Augustus Predictions** | **Glimmer Predictions** | **Braker3** |
| --- | --- | --- | --- | --- | --- |
| Core genes queried | 4896 | 4896 | 4896 | 4896 | 4896 |
| Complete | 4177 (85.31%) | 489 (9.99%) | 4552 (92.97%) | 4372 (89.30%) | 4769 (97.41%) |
| Complete + Partial | 4323 (88.30%) | 698 (14.26%) | 4645 (94.87%) | 4587 (93.69%) | 4781 (97.65%) |
| missing | 573 (11.70%) | 4198 (85.74%) | 251 (5.13%) | 309 (6.31%) | 115 (2.35%) |
| Average # of orthologs per core genes | 1.5 | 1.05 | 1.21 | 1.08 | 1.16 |
| % of detected core genes that have more than 1 ortholog | 30.33 | 4.29 | 19.24 | 6.91 | 13.23 |
| Scores in BUSCO format | C:85.3%[S:59.4%,D:25.9%],F:3.0%,M:11.7% | C:10.0%[S:9.6%,D:0.4%],F:4.3%,M:85.7%,n:4896 | C:93.0%[S:75.1%,D:17.9%],F:1.9%,M:5.1% | C:89.3%[S:83.1%,D:6.2%],F:4.4%,M:6.3% | C:97.4%[S:84.5%,D:12.9%],F:0.2%,M:2.4% |
| # of sequences | 35844 | 206825 | 302936 | 636815 | 35069 |
| Total length (nt) | 74170390 | 26945796 | 239588047 | 384961698 | 42311178 |
| Longest sequence (nt) | 28330 | 6299 | 46343 | 98788 | 16167 |
| Shortest sequence (nt) | 200 | 3 | 3 | 2 | 9 |
| Mean sequence length (nt) | 2069 | 130 | 791 | 605 | 1207 |
| Median sequence length (nt) | 1793 | 67 | 280 | 210 | 1011 |
| N50 sequence length (nt) | 2571 | 255 | 2520 | 1809 | 1551 |
| L50 sequence count | 9567 | 24822 | 24818 | 54231 | 8805 |

Table S7: SM gene pairs with Ka/Ks > 1

| **Gene 1** | **Gene 2** | **Ka** | **Ks** | **Ka/Ks** | **Annotation** |
| --- | --- | --- | --- | --- | --- |
| SM_01.g3515 | SM_05.g21252 | 0.549 | 0.248 | 2.21 | DOG1-like (seed dormancy) |
| SM_11.g6155 | SM_12.g8176 | 0.0148 | 0.0068 | 2.16 | Apyrase (ATP signaling) |
| SM_11.g6177 | SM_12.g8196 | 0.0447 | 0.0235 | 1.90 | Uncharacterized |
| SM_01.g3004 | SM_05.g21623 | 0.260 | 0.221 | 1.18 | Tyrosine decarboxylase |
| SM_11.g6236 | SM_12.g8262 | 0.0333 | 0.0300 | 1.11 | PGAM-like enzyme |

Table S8: Summary of major intragenomic synteny blocks in the Samba Mahsuri (SM) genome.
For each block, the corresponding chromosome pair, number of retained anchor gene pairs, mean synonymous divergence (Ks), estimated duplication age, and inferred evolutionary origin are shown. The majority of blocks trace back to the ancient σ-whole-genome duplication, whereas the Chr11–Chr12 block reflects a more recent segmental duplication.

| **Block** | **Chr Pair** | **Gene pairs** | **Mean Ks** | **Age (MYA)** | **Classification** |
| --- | --- | --- | --- | --- | --- |
| 27 | Chr1-Chr5 | 211 | 1.49 | ~115 | σ-WGD |
| 59 | Chr11-Chr12 | 114 | 0.40 | ~31 | Recent segmental duplication |
| 87 | Chr2-Chr4 | 164 | 1.59 | ~123 | σ-WGD |
| 93 | Chr2-Chr6 | 61 | 1.54 | ~118 | σ-WGD |
| 94 | Chr2-Chr6 | 35 | 1.39 | ~107 | σ-WGD |
